# Quantifying the effect of cereal plant trait plasticity on weed suppression in intercrops

**DOI:** 10.64898/2026.04.01.715874

**Authors:** David B. Kottelenberg, Alejandro Morales, Niels P. R. Anten, Lammert Bastiaans, Jochem B. Evers

## Abstract

In cereal-legume intercrops, weed suppression is primarily driven by cereals, whose competitiveness is shaped by trait plasticity—morphological adjustments in response to the intercrop environment. However, how individual cereal traits respond plastically and contribute to system performance remains unclear, hampering improvements through breeding or system design. We combined field experiments with functional-structural plant modelling to quantify plastic responses of four cereal traits (tiller number, tiller angle, specific leaf area (SLA), and specific internode length (SIL)) and their effects on weed suppression and crop productivity. Field measurements revealed plasticity in tiller number, tiller angle, and SIL between sole crops and intercrops, while SLA showed minimal differences. Simulations showed that intermediate tiller numbers resulted in the strongest weed suppression and highest productivity, indicating an optimum, while more horizontal tillers suppressed weeds slightly better than vertical ones. Weed suppression increased with higher SLA values, while SIL showed a saturating response, increasing to intermediate SIL values and plateauing thereafter. In simulations with short-statured cereal phenotypes (low SIL), the reduction in cereal weed suppression was compensated by the legume component. This study demonstrates how FSP modelling can be used to investigate trait plasticity mechanisms and generate testable hypotheses about trait effects in complex intercrop systems.

**Highlight:** Cereal trait plasticity shapes weed suppression in cereal-legume intercrops, with distinct response patterns per trait, while legumes can compensate for weakly competitive cereals, suggesting balanced competition over cereal dominance.

## 1 Introduction

Intercropping, the cultivation of multiple crop species on the same land, has been recognised as a means to enhance agricultural sustainability, with demonstrated benefits including enhanced yield stability and weed suppression (Wang et al., 2026; Gu et al., 2021; Li et al., 2020; Brooker et al., 2015; Bedoussac et al., 2014). The latter is particularly valuable given that weeds can significantly reduce crop yields (Oerke, 2006), while the widespread use of herbicides raises concerns about ecosystem functioning, human health, and the increasing prevalence of herbicide-resistant weed species (Arasan et al., 2025; Van Bruggen et al., 2018; Moss, 2017; Pérez et al., 2007).

Cereal-legume intercropping is one of the most prevalent intercropping strategies, combining the nitrogen-fixing ability of legumes with the strong competitive ability of cereals (Bybee-Finley and Ryan, 2018). As the typically dominant species, cereals drive weed suppression, often resulting in intercrops exceeding the average weed-suppressive ability of the component sole crops (Gu et al., 2021; Hauggaard-Nielsen et al., 2008). This dominance operates primarily through a selection effect, whereby the more competitive cereal captures a disproportionate share of resources, intensifying competitive asymmetry within the system (Gu et al., 2022). This is reflected in canopy cover and light capture dynamics that resemble those of the dominant cereal rather than the legume, as intercropped cereals cover the ground more efficiently and capture more light than when grown as sole crops (Kottelenberg et al., 2025; 2026a; Baumann et al., 2000). The magnitude of the selection effect depends on many aspects, including intercrop spatial arrangement, species choice, and environmental conditions (Salama et al., 2022; Wang et al., 2020; Stomph et al., 2020).

The increased competitive ability of cereals in intercrops is partly explained by plant trait plasticity, changes in plant traits in response to environmental conditions (Schneider, 2022). Here, we consider active and passive plasticity. Active plasticity occurs when a plant invokes physiological processes that alter its growth strategy in response to environmental signals. For example, as plants structures reflect and transmit most far-red light while absorbing most red light, plants perceive a reduced red:far-red (R:FR) light ratio as an indicator of future competition and respond by initiating a shade-avoidance response, which includes increased stem internode elongation and reduced tiller production (Franklin, 2008; Evers et al., 2006). Additionally, leaf traits such as declination angle and specific leaf area adapt plastically in intercropping environment, impacting light capture efficiency (Li et al., 2021; Zhu et al., 2015). Passive plasticity, on the other hand, occurs when plant traits change as an unavoidable consequence of physical or resource constraints rather than through a developmentally programmed response (Schneider, 2022). For instance, strong competition may limit available resources or growing space, resulting in reduced growth rates and smaller plant dimensions—not because the plant has actively adjusted its developmental strategy, but because environmental limitations physically constrain biomass accumulation or organ expansion.

The effects of active and passive plasticity are often difficult to disentangle. For example, increased competition may simultaneously trigger active shade-avoidance responses (altered R:FR ratios promoting elongation efficiency, i.e., more length per unit biomass invested) and passive constraints (reduced resource availability limiting overall growth). Disentangling how traits influence competition faces similar challenges: a change such as increased specific leaf area influences competitive outcomes through both direct geometric effects (intercepting light that would otherwise reach competitors) and whole-plant resource feedbacks (increased photosynthate production enabling greater overall growth and competitiveness across all traits). These effects operate through different pathways and can occur simultaneously, making it difficult to isolate the contribution of each mechanism to observed outcomes.

The effects of plasticity in individual plant traits on plant and crop performance are difficult to quantify in field experiments because multiple traits adapt simultaneously. Although field measurements can provide an estimate of the net effect of plasticity, they cannot isolate the contribution of individual trait contributions. Understanding which specific traits drive competitive outcomes can help to identify breeding targets and design effective intercropping systems. Functional-structural plant (FSP) models integrate mechanistic modelling of plant functions with 3D plant structure and topology, making them useful complementary tools for investigating plant-environment interactions and intra- and interspecific competition (Vos et al., 2010). FSP models have been shown to accurately simulate both sole crop and intercrop conditions and quantify the effect of individual plant traits on light capture by incorporating different plant phenotypes into diverse cropping scenarios (Li et al., 2021; Evers et al., 2019; Zhu et al., 2015). Additionally, FSP models have been established as tools for isolating individual trait effects on canopy development and light capture (Barillot et al., 2014; Evers et al., 2011; Sarlikioti et al., 2011; Kahlen et al., 2008; Evers et al., 2007), enabling systematic analysis of architectural trait contributions that are difficult to achieve through field experiments alone. Furthermore, FSP models enable investigation of trait interactions, whether traits act synergistically, additively, or antagonistically, which has important implications for breeding strategies that aim to combine multiple beneficial traits.

Field experiments have demonstrated that plant trait plasticity enhances weed suppression and increases crop yield in cereal-legume intercrops (MacLaren et al., 2023; Ajal et al., 2022). However, the extent to which individual plant traits adapt plastically in intercropping environments, and how these adaptations contribute to weed suppression and crop yield, is not well understood. Previous research has identified several cereal traits that respond to competitive environments and influence light capture: tillering (number and angle), which affects canopy structure and light distribution (Montazeaud et al., 2020; Zhu et al., 2015); specific internode length (SIL), which determines plant height and vertical positioning of leaves in the light gradient in the canopy (Li et al., 2010); and specific leaf area (SLA), which affects both leaf area deployment and photosynthetic capacity (Poorter et al., 2009). These traits are known to exhibit plasticity in response to shade signals and resource availability, making them candidates for understanding competitive mechanisms in intercrops. Understanding these mechanisms could inform breeding programmes and intercropping system design. In this paper, we therefore asked: which traits respond plastically to intercropping environments, through which mechanisms (active architectural positioning vs passive resource feedbacks) do these trait responses affect competition, and what are the effects on weed suppression and crop productivity? Given that cereals dominate cereal-legume intercrops and largely determine overall agronomic outcomes (Gu et al., 2021), this study focuses on cereal trait responses and their effects on weed control and crop productivity. To find which traits respond plastically and to what extent, we conducted four field experiments (2022-2024) measuring plant traits (tillering number, tiller angle, SIL, SLA) alongside weed biomass, crop biomass, and yield. Additionally, we developed and applied an FSP model of cereal-legume intercrops to quantify how individual trait plasticity contributes to system performance. Based on our model we developed a method to explore the extent to which observed plastic responses can considered to be active or passive.

## 2 Materials and Methods

### 2.1 Field experiments

In this study, we used data from the 2022, 2023-SP, 2023-RD, and 2024 field experiments as described in Kottelenberg et al. (2026a). We conducted these experiments in Wageningen, the Netherlands (51° 59’20.4”N 5° 39’07.0”E). Daily temperature average and rainfall during the experiments can be found in Supplementary Fig. S.1. In all experiments, the soil was of a loamy sand type, with approximately 3% organic matter, 3% clay, 15% silt, and 79% sand. The 2022 and 2023 experimental fields had previously been cultivated with winter wheat while the 2024 experimental field had previously been cultivated with sugar beet. All experiments used a randomised complete block design.

In 2022, we tested four cereal species: barley (*Hordeum vulgare* cv. Irina), rye (*Secale cereale* cv. Boyce), triticale (× *Triticosecale* cv. Santos), and wheat (*Triticum aestivum* cv. Quintus), combined with three legume species: pea (*Pisum arvense* cv. Tiberius), lupine (*Lupinus albus* cv. Frieda), and faba bean (*Vicia faba* cv. Cartouche). These were grown as sole crops and all possible cereal-legume combinations as 1:1 alternating row intercrops with 12.5 cm row spacing (Supplementary Fig. S.2). Plot size was 3 × 11 m with four replicates. Based on these results, triticale-faba bean was selected for subsequent experiments.

In 2023, two experiments examined specific design factors using triticale and faba bean. The first experiment (2023-SP) tested effects of species proportions and density. Triticale (cv. Mazur) and faba bean (cv. Cartouche) were grown as sole crops at standard density (T, F) and triticale at 1.5× density (T+), alongside intercrops with varying row ratios: 1:1 alternating rows (1T:1F), 3:1 rows (3T:1F), 1:3 rows (1T:3F), and 1:1 within-row mixed (1T:1F-M), all at 12.5 cm row spacing (Supplementary Fig. S.3). Plot size was 3 × 7 m with five replicates. The second experiment (2023-RD) tested effects of row distance and spatial arrangement.

Triticale (cv. Mazur) and faba bean (cv. Cartouche) were grown as sole crops (T, F) and as 1:1 intercrops, either row-alternating (1T:1F) or within-row mixed (1T:1F-M), at two row distances: 12.5 cm (T, F, 1T:1F, 1T:1F-M) and 37.5 cm (T-375, F-375, 1T:1F-375, 1T:1F-M-375) (Supplementary Fig. S.3). Plot size was 3 × 16 m with four replicates.

In 2024, triticale (cv. Dublet) and faba bean (cv. Cartouche) were grown as sole crops (T, F) and as 1:1 row-alternating intercrop (1T:1F) at 12.5 cm row spacing (Supplementary Fig. S.4). A germination test showed approximately 90% triticale seed viability compared to nearly 100% in previous years, and sowing rates were adjusted accordingly. Additionally, faba bean seeding rate was increased based on weak weed suppression, sparse canopies at lower densities, and updated agronomic recommendations (Supplementary Table S.1). Plot size was 3 × 11 m with five replicates.

Sowing dates were 19 April 2022, 2 March 2023, and 12 April 2024, using the Macon trial field sowing machine. Seeding rates and densities can be found in Supplementary Table S.1. All intercrops were sown as replacement designs, maintaining the same per-area density as corresponding sole crops.

To investigate plant trait plasticity in intercrops without weed interference, plots were treated with herbicides. In 2022, plots were treated with 2 L haL¹ Basagran (BASF, 480 g LL¹ bentazon) on 12 May. In 2023, plots were treated with 2 L haL¹ soil herbicide Stomp (BASF, 400 g LL¹ pendimethalin) on 28 February, two days before sowing. In 2024, plots were treated with 1 L haL¹ Stomp on 11 April, a day before sowing, and 2 L haL¹ Basagran on 15 May.

In 2022, plots containing triticale received treatments against powdery mildew with 0.5 L haL¹ Nissodium (Certis Belchim, 50 g LL¹ cyflufenamid) and 2 L haL¹ manganese nitrate on 14 June. In 2023, plots containing faba bean were treated with 1 L haL¹ Prosaro (Bayer, 125 g LL¹ prothioconazole and 125 g LL¹ tebuconazole) on 20 June to combat chocolate spot disease (Botrytis fabae). Irrigation was applied during periods of drought on 16 June and 21 June 2023, where approximately 15 mm of groundwater was applied per date. Prior to sowing, the soil was ploughed, fertilised, and harrowed. All plots received a pre-sowing base fertilisation of 50 kg haL¹ PLOL and 120 kg haL¹ KLO. Additionally, post-sowing fertilisation was done with 296 kg haL¹ calcium ammonium nitrate (CAN; 80 kg N haL¹) for triticale and 111 kg haL¹ CAN (30 kg N haL¹) for faba bean, with proportional adjustments for intercrops.

### 2.2 Crop measurements

Cereal tillering was measured weekly between 25 May and 14 June in 2022, 17 April and 31 May in 2023, and 10 May and 13 June in 2024. Measurements were done by counting the number of stems along 30 cm of cereal row, replicated five times per plot. Cereal tiller angle was measured on 21 June 2022 and 23 May 2023 by photographing cereal plants along the row direction, with tillers visible extending laterally toward adjacent rows, with two adjacent rows visible per image. For the first visible plant in each of the two rows (two plants per plot), the angle of two randomly selected tillers was measured from vertical using ImageJ, with 0° representing upright growth and 90° representing horizontal growth. Internode and leaf data was collected on 12, 18, and 26 April and 3, 12, 19, and 25 May 2023 by taking three random plants from designated destructive harvest areas in the field (Supplementary Fig. S.3). Leaves and internodes were separated from the top three phytomers (or fewer, if fewer than three phytomers were present) of the three plants. Leaf area was measured using the LI-COR LI-3100C leaf area scanner, and internode length and width (in cm) were measured. The internodes and leaves were dried at 105 °C for 24 hours, after which the dry weight was taken. SLA was calculated as the leaf area per unit biomass of the leaf in cm² mgL¹, and SIL was calculated as the internode length per unit biomass of the internode in mm mgL¹. Internodes and leaves were separately measured by phytomer rank. Internode data was only available on the last three measurement dates, as internode elongation had not started yet on the earlier dates.

### 2.3 Functional-structural plant model

#### 2.3.1 Model description

The FSP model was developed using the virtual plant lab (VPL) modelling platform in the Julia programming language (Morales et al., 2025; www.virtualplantlab.com). The model simulates plant growth and development by integrating physiological processes with 3D plant architecture, enabling mechanistic simulation of light competition and assimilate allocation. Species-specific traits are represented through parameter sets, while the underlying model structure remains constant across species.

Light interception is calculated using a stochastic forward ray tracing algorithm that tracks radiation through the 3D canopy structure, using an overcast sky light regime that combines diffuse and direct light based on Spitters et al. (1986). At each daily timestep, the model computes photosynthetically active radiation (PAR) absorbed by individual leaves based on their position, orientation, and shading by surrounding vegetation. To capture diurnal variation in light angles, PAR absorption is calculated at ten time points between sunrise and sunset using Gaussian quadrature integration. The ray tracer simulates light in three spectral channels—PAR, red, and far-red—each with distinct optical properties when interacting with canopy structures. Leaves and stems can have different reflectance and transmittance values for red and far-red wavelengths (see species parameters in Supplementary Tables S.2-S.4). If parameters values are set such that that red light is being mostly absorbed and far-red light being mostly reflected and transmitted, this differential interaction causes R:FR ratios of the light inside canopies to decline as the plants grow and develop, providing environmental signals that trigger shade avoidance responses such as increased stem elongation and reduced tillering (see section 2.3.2 below).

Plant development proceeds through sequential phytomer production, with each phytomer consisting of a node, internode, leaf, and a branch bud. Phytomer appearance rate is temperature-driven, following thermal time accumulation. Organ growth parameters are genetic but respond plastically to environmental signals including R:FR ratios and source:sink dynamics, enabling the model to simulate both active and passive plastic responses to competitive environments.

Carbon assimilation and allocation follow a source-sink framework where photosynthesis generates assimilates (source) that are allocated to growing phytomer organs (sinks) (Evers et al., 2010). Daily gross photosynthesis is calculated for each leaf and internode using the Farquhar-von Caemmerer-Berry (FvCB) C3 photosynthesis model, with key biochemical parameters derived from leaf nitrogen content following empirical relationships established by Yin and Struik (2009) (Vcmax25: maximum rate of carboxylation at 25 °C (µmol mol^-1^); Jmax25: maximum rate of electron transport at 25 °C (µmol m^-2^ s^-1^); TPU25: maximum rate of triose phosphate utilisation at 25 °C (µmol m^-2^ s^-1^)). Photosynthesis rates depend on absorbed PAR and environmental conditions, with respiration costs subtracted to determine net assimilate production. The source:sink ratio, calculated as available assimilates divided by total sink demand, regulates growth processes including leaf expansion and internode elongation on main stem and branches. When the plant source:sink ratio falls below 1.0, growth is reduced proportionally across all sinks, i.e., no sink hierarchy is assumed. Each organ has a defined growth duration during which it acts as a sink. The sink strength of growing organs follows a parabolic curve over time, calculated from a potential biomass parameter and the organ’s age relative to a maximum growth age (set at half the total growth duration), such that sink strength increases early in development, peaks at the midpoint, then declines (Evers et al., 2010).

Based on Cao et al. (2022), leaf angle (defined here as the declination angle relative to vertical) was different depending on the phytomer number on the plant stem, with leaves higher on the plant having a smaller angle with respect to the stem (i.e., being more vertically inclined). This is based on a logistic function:

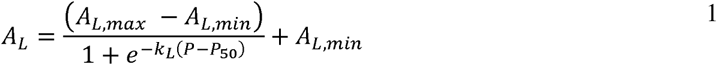

where A_L_ is the leaf angle, A_L,max_ and A_L,min_ are the maximum and minimum leaf angles, respectively, k_L_ is the initial rate of increase of the leaf angle per phytomer number, P is the phytomer number, and P_50_ is the phytomer number at which the leaf angle is halfway between its maximum and minimum value.

Finally, a cereal and legume species were modelled using parameters based on available data on triticale and faba bean sole crops from the 2023-SP experiment in Kottelenberg et al., 2026a, and this study. Model parameters that we could not measure were estimated from literature. For cereals, 12 parameters were directly measured or calibrated from our field experiments (tiller number, tiller angle, SIL, SLA, and key phenological and organ growth parameters), while 25 were taken from literature on triticale or related cereals (primarily photosynthetic, respiratory, and optical properties) (Supplementary Table S.2). For legumes, 11 parameters came from field measurements and 18 from literature sources (Supplementary Table S.3). As such, the modelled cereals and legumes roughly represent our experimental species triticale and faba bean.

A generic broadleaf weed species was modelled using parameters based on *Chenopodium quinoa* Willd. from the FSPM_BASIC^1^ model, as this species is morphologically very similar to its congener *Chenopodium album* L., by far the most common weed species in our experimental fields (Supplementary Table S.4). To represent intraspecific variation within the weed population, specific internode length and specific leaf area values were randomly drawn from uniform distributions per plant (SIL: 0.1–0.6 mm mgL¹; SLA: 0.17 and 1.0 cm^2^ mg^-1^). This generated a range of weed phenotypes with varying competitive abilities, representing natural trait variation in weed populations where individual plants differ in growth strategy (e.g., fast-growing with high SLA versus slower-growing with low SLA), potentially influencing competitive outcomes with crop species. Weeds were modelled with shade avoidance responses for internode elongation (R:FR-mediated, similar to crop species, see section 2.3.2) but without plasticity in other traits (branch angle, SLA, or branch production), representing a simplified weed competitor with fixed architectural characteristics but responsive stem elongation.

At model initialisation, each plant is assigned a random azimuthal orientation (0–360°), resulting in variation in orientation of plants. Additionally, the ray tracing light model is stochastic, introducing variation in light conditions between replicate simulations.

#### 2.3.2 Plant trait plasticity

We implemented plastic responses of the crop species in four traits in the model: tiller number, tiller angle, elongation (through SIL), and SLA (Fig. 1a).

**Figure 1.**
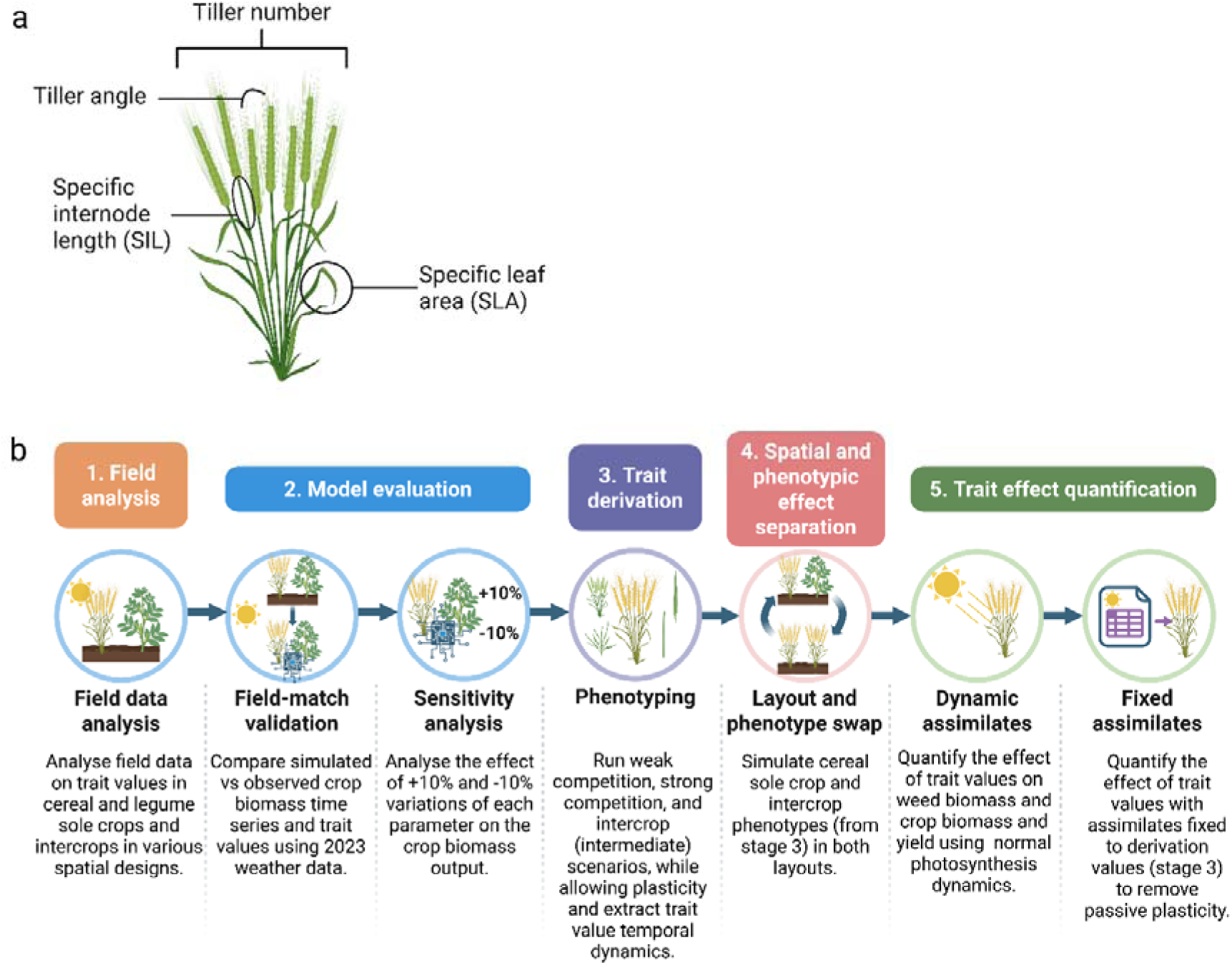
Focal cereal traits and workflow. (a) Schematic representation of a cereal plant indicating the four morphological traits measured in field experiments and manipulated in simulations: tiller number, tiller declination angle (°), specific internode length (SIL, mm mgL¹), and specific leaf area (SLA, cm² mgL¹). (b) Integrated field-model workflow with five stages: (1) field experiments, measuring cereal trait responses across sole crops and intercrops to quantify plasticity and parameterise the model; (2) model evaluation, comparing simulated outputs against field measurements and assessing parameter sensitivity; (3) trait derivation, extracting trait values across a competition gradient; (4) spatial and phenotypic effect separation, isolating effects of spatial arrangement from trait plasticity; and (5) trait effect quantification, determining individual trait effects on system performance under dynamic and fixed assimilation modes.

The parameters governing each plasticity function were calibrated to field measurements and kept constant across all simulations, representing a single genotype. The different trait values produced across scenarios (section 2.3.3, stage 3) therefore represent phenotypic responses of this genotype to different competitive environments, not genotypic variation.

In the model, cereal tiller number can either be fixed or respond plastically to the local light environment. When tiller number is fixed, a predetermined number of tillers is initiated regardless of competition. When plastic tillering is enabled, tiller production is controlled through an R:FR threshold mechanism based on R:FR ratios detected at the plant base. Tillering is blocked until plants reach a specified minimum age. Subsequently, at each daily timestep, the oldest dormant branch bud initiates and begins growing when the R:FR ratio at the base exceeds a threshold value. A branch delay parameter enforces a minimum number of days between successive bud initiations, reflecting the tiller emergence rate measured in the field. Tiller production ceases when either the R:FR ratio falls below the threshold (indicating competition from neighbouring plants; reversible, e.g. when neighbouring plants are removed) or when flowering starts (irreversible). The R:FR threshold value controls the sensitivity of tiller number to competition: values close to the unfiltered sunlight ratio of 1.2 cause tillering to cease as soon as any neighbouring vegetation is detected, while values close to 0.0 result in near-maximum tiller numbers regardless of competition. Intermediate threshold values produce plastic tiller numbers that depend on the competitive environment. Additionally, branch abortion follows Larue et al. (2019): branches are aborted when their source-to-maintenance ratio (assimilates produced via photosynthesis of the branch’s leaves divided by maintenance respiration costs of leaf and branch tissue) remains below a threshold for a specified duration.

Legume plants were modelled without branching because minimal branching was observed in the field even at the lowest competition level, and only scenarios with equal or greater legume competition are simulated. Furthermore, the model focused on cereal trait plasticity rather than legume architectural responses.

Tiller angle can either be fixed or respond plastically to the local light environment. When fixed, all tillers are assigned a constant angle regardless of competition. When plastic, tiller angle is modelled using a power-law response function based on the R:FR ratio detected at the plant base:

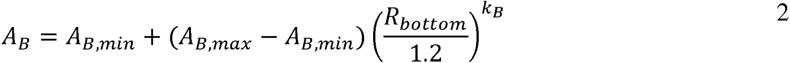

where A_B_ is the tiller angle (degrees from vertical), A_B,max_ and A_B,min_ are the maximum and minimum tiller angles, k_B_ is the shape parameter controlling response steepness, and R_bottom_ is the red:far-red ratio detected by a sensor at the base of the plant stem. R_bottom_ represents the light environment experienced at ground level, where low R:FR values indicate competition from neighbouring plants. The range of plasticity is therefore bounded by A_B,max_ and A_B,min_, which define the tiller angles under minimum and maximum competition, respectively, while k_B_ controls how sharply the transition between these extremes occurs. A_B,max_, A_B,min_, and k_B_ were calibrated to field measurements (Supplementary Table S.2).

The denominator 1.2 corresponds to the R:FR ratio of incoming sunlight, so that 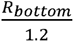 approaches 1.0 in the absence of canopy filtering and decreases towards 0 under dense vegetation.

Tiller angle is the declination angle between the main stem and the first internode of the tiller, determined at the time of tiller initiation based on the R_bottom_ value of that moment. This initial angle remains fixed throughout that tiller’s development, though subsequent internodes on the same tiller progressively orient more vertically due to gravitropism. Consequently, tillers emerging at different times can have different angles depending on the prevailing R:FR conditions. This represents the developmental plasticity observed in cereal tillering, where tiller angle is established during early development rather than continuously adjusted in mature tillers.

SLA can either be fixed or respond plastically to the local light environment. When fixed, all leaves are assigned a constant SLA regardless of light capture. When plastic, SLA is modelled using a logistic function based on the leaf’s capture of photosynthetically active radiation (PAR), following Poorter et al. (2009):

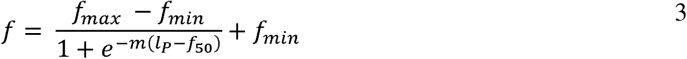

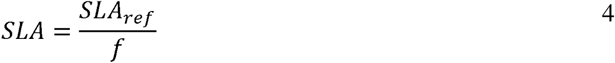

where *f* is the plasticity factor, *f_max_* and *f_min_* are the maximum and minimum value of the SLA plasticity factor, *m* is the shape parameter defining the steepness of the sigmoid curve, *l_p_* is the PAR absorbed by the leaf (µmol m^-2^), *f_50_* is the *l_p_* -value where *f* equals the midpoint between *f_max_* and *f_min_*, SLA is the specific leaf area (cm mg ), and *SLA_ref_* is the reference SLA at *f* = 1. The parameters *f_max_* , *f_min_*, *m*, *f*_50_, were based on the relationship found by Poorter et al. (2009) and *SLA_ref_* was calibrated to field measurements (Supplementary Table S.2). The range of plasticity is bounded by *f_max_* and *f_min_*, while m and *f*_50_ control the shape of the response to light capture.

SLA is calculated per leaf and per timestep, so that leaves developing at different times and canopy positions differ in leaf area per unit biomass. A higher SLA increases the leaf area produced per unit biomass investment, but reduces biomass per area and consequently nitrogen content per area, which decreases the per-area photosynthesis rate.

SIL can either be fixed or respond plastically to the local light environment. When fixed, all internodes elongate with a constant length per unit biomass regardless of competition. When plastic, the SIL is modulated by an elongation factor (E), determined by the R:FR ratio of light captured by a 1 cm sensor around the apex of the main stem:

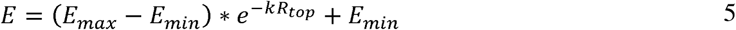

where *E_max_* is the maximum elongation factor, *E_min_* is the minimum elongation factor, *k* is the shape parameter controlling response sensitivity, and *R_top_* is the red:far-red ratio detected by the apex sensor. *R_top_* represents the light environment at the growing point, where low R:FR values indicate shading from neighbouring plants, triggering the shade avoidance elongation response. The range of plasticity is bounded by *E_max_* and *E_min_*, while *k* controls how sharply the elongation response changes with R:FR. The parameters *E_max_* , *E_min_*, and *k* were calibrated to field measurements (Supplementary Tables S.2-S.4).

The elongation factor is calculated daily and applied to all actively elongating internodes. Each internode elongates during its active growth period, accumulating length according to the daily *E* value and available assimilates. Internodes that have completed their growth period no longer respond to changes in *R_top_*, so that shade signals at the apex influence elongation efficiency only during active growth.

#### 2.3.3 Simulation setup

To quantify how plant trait plasticity affects weed suppression and crop productivity, we followed a five-stage workflow (Fig. 1b). Stage 1, field data analysis, is described above. The remaining four stages constitute the simulation experiment: model evaluation (stage 2), trait derivation (stage 3), spatial and phenotypic effect separation (stage 4), and trait effect quantification (stage 5).

While the model was parameterised based on sole crop data, in the second stage, model evaluation, we validated the model and performed a sensitivity analysis on 1:1 ratio cereal-legume row intercrops (1T:1F), all without weeds, matching part of the setup of the 2023 field experiment. We used observed radiation and temperature data from the 2023 growing season as model inputs (KNMI, 2025) and compared simulated biomass, yield, tiller number, and tiller angle (triticale only) against field measurements of the same treatments on the field measurement dates.

We assessed parameter sensitivity by performing a local sensitivity analysis on the 1:1 intercropping system. For each numeric input parameter for both cereal and legume (72 total), we conducted separate simulations where that parameter was increased or decreased by 10% while all other parameters remained at their default values (144 unique simulations). This was done for three replicates (432 total simulations). We then compared the resulting mean above-ground biomass and crop yield to baseline values from the default parameterisation, averaged over the three replicate runs for each parameter variation. This one-at-a-time approach isolated the individual contribution of each parameter to model predictions (Supplementary Fig. S.5).

The third stage characterised cereal trait responses across a competition gradient. We simulated five cropping scenarios representing increasing competitive intensity: (1) isolated cereal plants (no competition), (2) half-density cereal-legume intercrops without weeds (weak competition), (3) cereal-legume intercrops with weeds (intermediate competition), (4) cereal sole crop with weeds (strong competition), and (5) double-density cereal sole crops with weeds (extreme competition). In each scenario, the plasticity functions described in section 2.3.2 were enabled for all four traits, so that tiller number, tiller angle, SLA, and SIL emerged from the model’s responses to the competitive environment. Simulations used long-term average radiation and temperature data (2000–2024) to remove year-to-year environmental noise. We acknowledge that the extreme scenarios (isolated plants, half-density intercrops without weeds, and double-density sole crops) were not validated against field data, so the simulated trait trajectories identify trends in plastic responses rather than quantitatively validated predictions. From these simulations, we extracted mean trait values at every daily timestep. These extracted values were used to parameterise subsequent trait manipulation experiments: trait values from the weedy sole crop and intercrop scenarios were used in stage four, while values from all five scenarios provided focal points spanning the full competition range for stage five.

In the fourth stage, spatial and phenotypic effect separation, we isolated the effects of spatial arrangement from trait plasticity (comparable to Li et al., 2021; Zhu et al., 2015) through virtual transplant simulations. We conducted four simulation scenarios combining two spatial arrangements (weed-infested cereal sole crop vs. weed-infested cereal-legume intercrop) with two sets of cereal trait values (the four trait trajectories from stage three: tiller number, tiller angle, SLA, and SIL from either the sole crop scenario or the intercrop scenario). This created a 2×2 factorial design: (1) sole crop spatial arrangement with sole crop trait values, (2) sole crop spatial arrangement with intercrop trait values, (3) intercrop spatial arrangement with intercrop trait values, and (4) intercrop spatial arrangement with sole crop trait values. Weeds were placed randomly in the field area across all scenarios. This design separated spatial effects (row arrangement, neighbour identity) from trait plasticity effects (differences in the four trait trajectories between sole crops and intercrops), allowing us to quantify whether performance differences arose from plastic trait adjustments or from spatial arrangement itself.

In the fifth stage, trait effect quantification, we quantified how individual trait variation affects system performance by simulating intercrops where three traits followed their intermediate-competition dynamics (from the weedy cereal-legume intercrop scenario in the third stage) while one focal trait was set to either no-competition dynamics (isolated plants scenario), weak-competition dynamics (half-density intercrop scenario), strong competition dynamics (weed-infested cereal sole crop scenario), or extreme-competition dynamics (double-density sole crops scenario). Although traits varied temporally according to their developmental patterns (tiller number through production and abortion, tiller angle through development, SIL through R:FR dynamics, SLA through PAR-mediated dynamics), these temporal patterns were predetermined inputs from the third stage. This factorial approach generated 16 simulation scenarios: no-competition, weak-competition, strong-competition, and extreme-competition variants for each of the four traits (Table 1).

**Table 1.** Factorial design for trait effect quantification simulations. Each row represents one simulation where one focal trait trajectory was set to no-competition dynamics (isolated plants; ”No competition”), weak-competition dynamics (half-density weed-free intercrop; ”Weak”), strong-competition dynamics (weed-infested cereal sole crops; ”Strong”), or strong-competition dynamics (double-density weed-infested sole crops; ”Extreme”), while the other three trait trajectories followed intermediate-competition (weed-infested intercrop; ”Intermediate”) dynamics. This created sixteen simulations: four competition levels for each of the four traits. All simulations were conducted under both dynamic assimilation (where cereal light capture determines available assimilates, creating resource feedbacks) and fixed assimilation (where total cereal assimilates per plant were held constant at values from the intermediate-competition scenario in stage three, eliminating whole-plant resource feedbacks while maintaining within-plant source-sink dynamics) modes.

| Simulation | Tiller number | Tiller angle | SLA | SIL |
| --- | --- | --- | --- | --- |
| 1 | No competition | Intermediate | Intermediate | Intermediate |
| 2 | Weak | Intermediate | Intermediate | Intermediate |
| 3 | Strong | Intermediate | Intermediate | Intermediate |
| 4 | Extreme | Intermediate | Intermediate | Intermediate |
| 5 | Intermediate | No competition | Intermediate | Intermediate |
| 6 | Intermediate | Weak | Intermediate | Intermediate |
| 7 | Intermediate | Strong | Intermediate | Intermediate |
| 8 | Intermediate | Extreme | Intermediate | Intermediate |
| 9 | Intermediate | Intermediate | No competition | Intermediate |
| 10 | Intermediate | Intermediate | Weak | Intermediate |
| 11 | Intermediate | Intermediate | Strong | Intermediate |
| 12 | Intermediate | Intermediate | Extreme | Intermediate |
| 13 | Intermediate | Intermediate | Intermediate | No competition |
| 14 | Intermediate | Intermediate | Intermediate | Weak |
| 15 | Intermediate | Intermediate | Intermediate | Strong |
| 16 | Intermediate | Intermediate | Intermediate | Extreme |

To distinguish active plasticity (developmental responses to environmental signals) from passive plasticity (resource-limitation effects), we conducted the fourth stage simulations under two assimilation modes. Under dynamic assimilation (default), cereal photosynthetic assimilates were calculated from simulated light capture, allowing resource feedbacks to operate (e.g., the high-SLA phenotype has a larger leaf area, potentially capturing more light, resulting in even more leaf area), and therefore representing both active and passive plasticity. Under fixed assimilation, cereal assimilates were held constant at values from the intermediate-competition intercrop simulation (from the third stage), isolating geometric and allocation effects while eliminating resource feedbacks, thereby representing the effects of active plasticity without passive responses. Legume and weed assimilates remained dynamic in all simulations.

Under fixed trait values and assimilation, imposing uniform trait values or resource capture dynamics across all plants removed the adaptive capacity present in the original parameterisation simulations, where individual plants adapted to local competitive environments. This caused the intercrop phenotype to perform sub-optimally under fixed assimilation. Therefore, we compared patterns within each assimilation mode separately rather than between modes.

### 2.4 Statistical analysis

We tested whether tiller number, tiller angle, SLA, and SIL of cereals differed among cropping treatments in the field experiment using a linear-mixed-effects model. The model included cropping treatment as a fixed effect and block (replicate) as a random effect. Model residuals were assessed for normality using the Shapiro–Wilk test and for homoscedasticity with Levene’s test. If these assumptions were violated, a Box–Cox transformation was applied to the response variable data (Box and Cox, 1964), which subsequently always satisfied the assumptions. Statistical significance of treatment effects was assessed using analysis of variance with Satterthwaite’s approximation for denominator degrees of freedom (lmerTest package in R). When a significant effect was detected (P < 0.05), post-hoc pairwise comparisons were performed using Tukey-adjusted estimated marginal means, and groupings of variable means were visualised using compact letter displays. All analyses were conducted in R 4.5.0 (R Core Team 2025).

## 3. Results

### 3.1 Field analysis

In the first stage of the workflow we analysed cereal trait plasticity in the field experiments (Fig. 1b, stage 1). Plasticity in final tiller number was apparent in some treatments. Barley and triticale had more tillers in intercrops with faba bean than in sole crops (Fig. 2a, d). Furthermore, tiller number increased as the triticale proportion in the intercrop decreased (Fig. 2b). Most apparent was the large increase in number of tillers in the 37.5 cm row-distance crops compared to the 12.5 cm row distance (Fig. 2c). This increase was less in the 37.5 cm intercrop where the plants were mixed in the rows (1T:1F-M-375).

**Figure 2.**
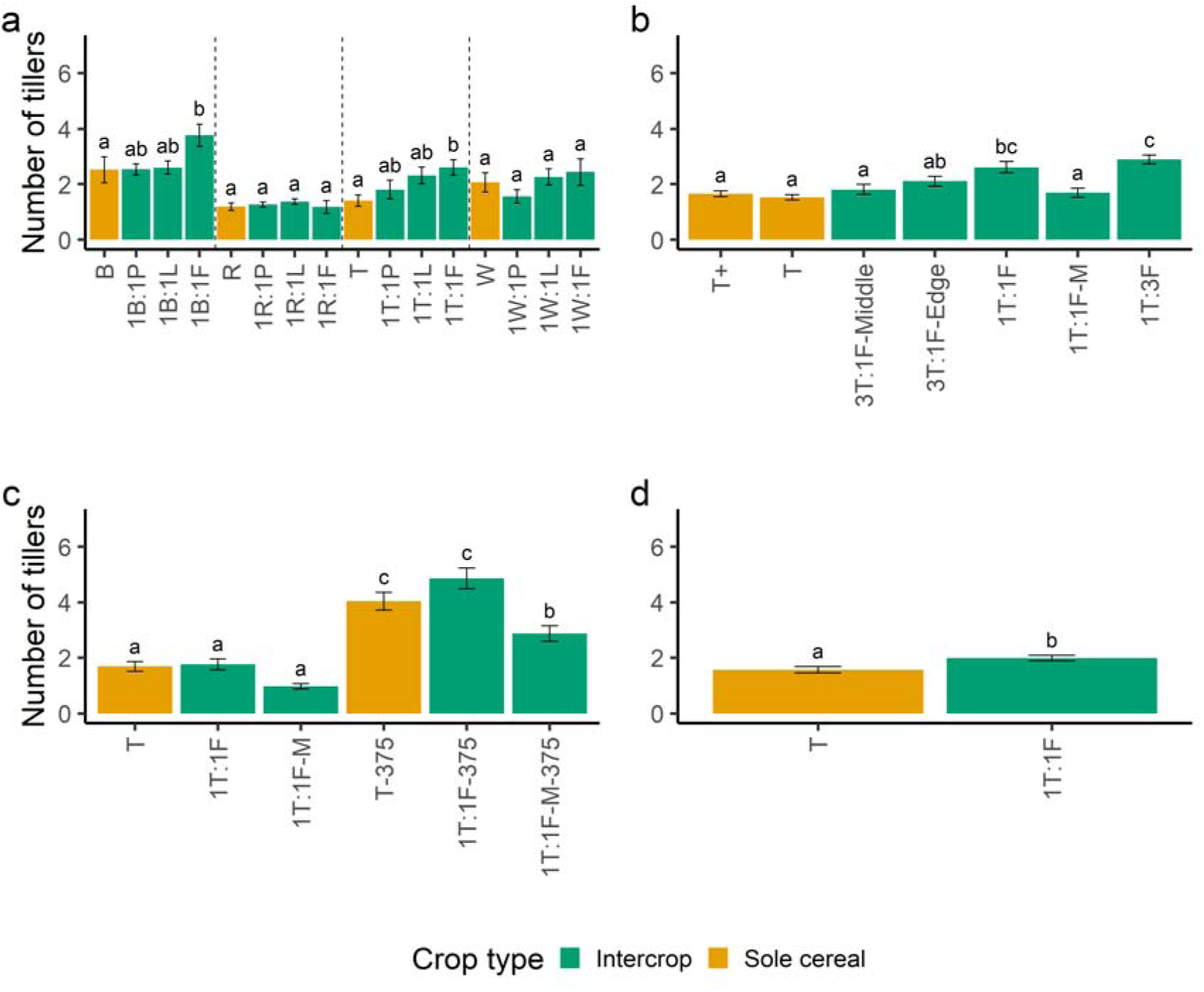
Cereal tiller number. Final number of tillers (excluding the main stem) per cereal plant in sole crops (red) and intercrops (green) across treatments. Treatments in the 2022 experiment (a) comprised sole crops of barley (B), rye (R), triticale (T), wheat (W), pea (P), lupine (L), and faba bean (F), together with cereal–legume row intercrops at a 1:1 ratio (1B:1P, etc.). Treatments in the 2023-SP experiment (b) comprised triticale sole crops at standard density (T) and at 1.5 times that density (T+), and intercrops of triticale and faba bean arranged as 1:1 row ratio (1T:1F), mixed within rows (1T:1F-M), and row ratios of 1:3 (1T:3F) and 3:1 (3T:1F). In the 3T:1F treatment, ‘Middle’ and ‘Edge’ indicate the middle row and edge rows of the 3-row strip. The 2023-RD experiment included T, 1T:1F, and 1T:1F-M similar to the 2023-SP experiment, as well as these treatments with 37.5 cm row spacing (−375). Treatments in the 2024 experiment (d) included T and 1T:1F. Error bars indicate the standard error. Different letters indicate significant differences between treatments at P < 0.05; for 2022 the letters are grouped per cereal species, e.g., a comparison is made for the barley group between barley, barley-pea, barley-lupine, and barley-faba, similarly for the other cereals.

The angle of cereal tillers with respect to the main stem showed differences between sole crop and intercrop in rye and triticale systems, where intercropped cereals had larger tiller angles (tillers more horizontally projected), indicating wider tillering and thereby occupying more space (Fig. 3a). Furthermore, wider tillers were seen at lower triticale proportions (Fig. 3b) and larger row distances (Fib. 3c). The within-row mixing at larger row distance (1T:1F-M-375) showed an intermediate tiller angle between the 12.5 cm row distance systems and the non-mixed 37.5 cm row distance systems.

**Figure 3.**
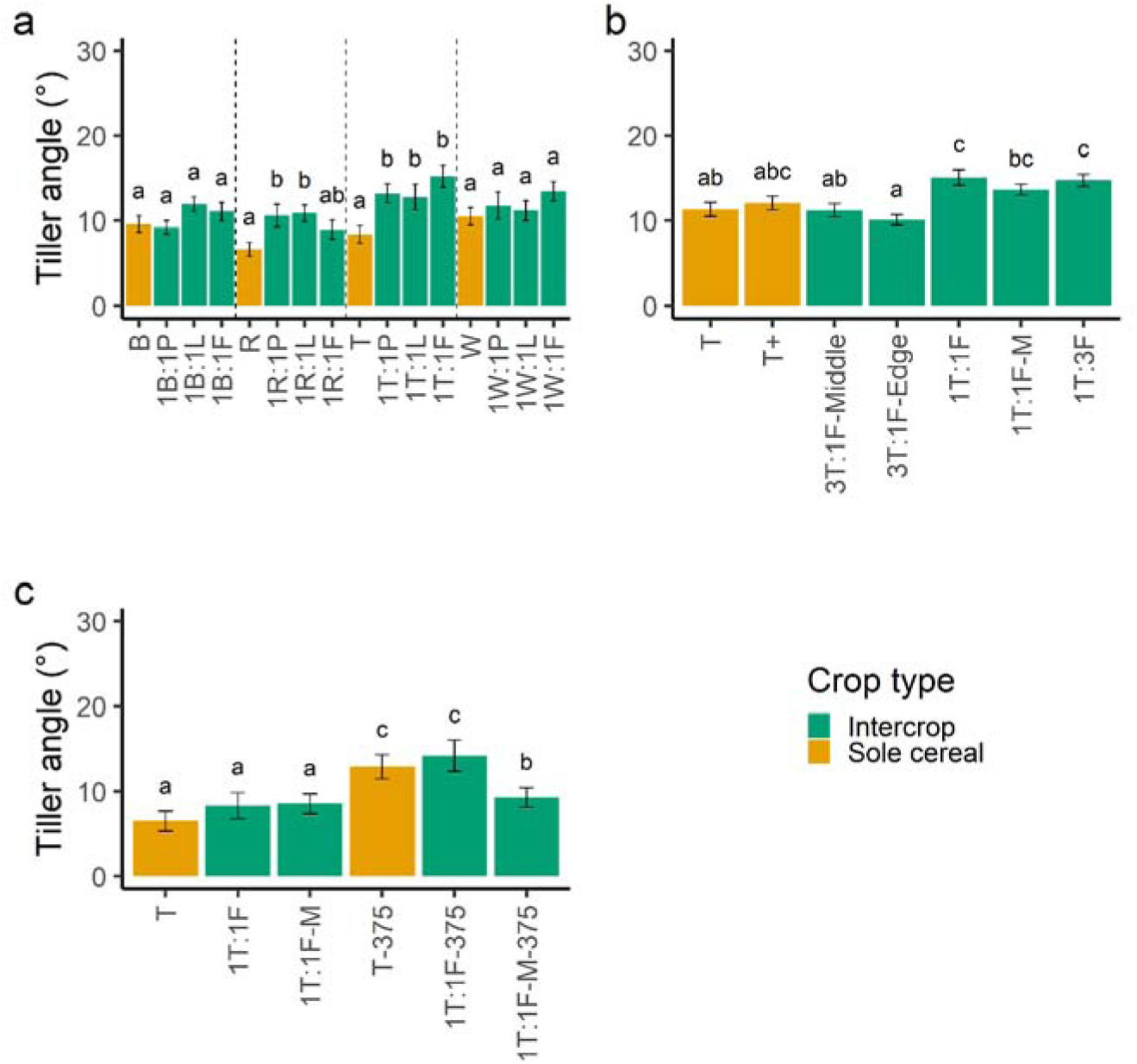
Cereal tiller declination angle. The angle of cereal tillers with respect to the main stem (larger values indicate more horizontal tillers) in sole crops (orange) and intercrops (green) across treatments for the (a) 2022, (b) 2023-SP, (c) 2023-RD, and (d) 2024 experiments. Treatments in the 2022 experiment (a) comprised sole crops of barley (B), rye (R), triticale (T), wheat (W), pea (P), lupine (L), and faba bean (F), together with cereal–legume row intercrops at a 1:1 ratio (1B:1P, etc.). Treatments in the 2023-SP experiment (b) comprised of triticale sole crops at standard density (T) and at 1.5 times that density (T+), and intercrops of triticale and faba bean arranged as 1:1 row ratio (1T:1F), mixed within rows (1T:1F-M), and row ratios of 1:3 (1T:3F) and 3:1 (3T:1F). In the 3T:1F treatment, ‘Middle’ and ‘Edge’ indicate the middle row and edge rows of the 3-row strip. The 2023-RD experiment (c) included T, 1T:1F, and 1T:1F-M similar to the 2023-SP experiment, as well as these treatments with 37.5 cm row spacing (−375). Error bars indicate standard errors. Different letters indicate significant differences between treatments at P < 0.05; for 2022 the letters are grouped per cereal species, e.g., a comparison is made for the barley group between barley, barley-pea, barley-lupine, and barley-faba, similarly for the other cereals.

No significant treatment effects on the SLA were detected in the 2023 experiment (Fig. 4). The SLAs showed a similar range of values between ranks as well, indicating that the SLA did not change much between sole crop or intercrop environments or location on the stem when comparing the first three ranks.

**Figure 4.**
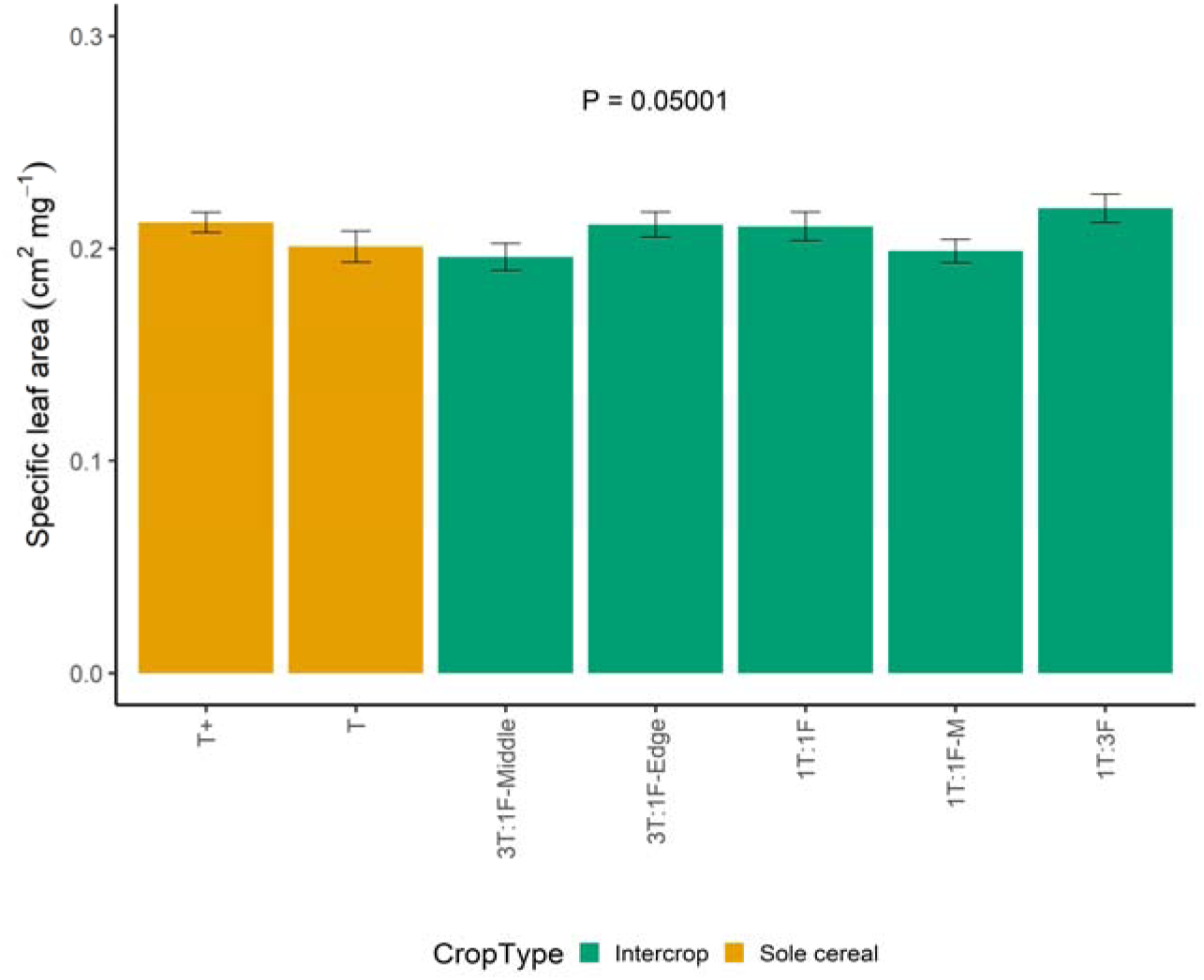
Cereal specific leaf area. Specific leaf area (SLA; in cm^2^ mg^-1^) of cereal plants in the 2023-SP sole crops and intercrops. Treatments include sole crop at standard density (T), sole crops at 1.5 times the standard density (T+), and intercrops with triticale and faba bean at a 1:1 row ratio (1T:1F), mixed within the row (1T:1F-M), and at row ratios of 1:3 (1T:3F) and 3:1 (3T:1F). ‘Middle’ and ‘Edge’ indicate the middle row and edge rows of the 3-row strip in the 3T:1F treatment. Error bars indicate the standard error. The P-value displayed is from an ANOVA (F-test with Satterthwaite’s approximation) comparing treatments.

SIL showed minimal plasticity between sole crops and intercrops across most treatments and phytomer positions (Fig. 5). SIL values increased from the first internode to the third internode, reflecting developmental changes in internode structure along the stem. However, within each phytomer position, differences between sole crops and intercrops were small, though a pattern suggests that treatments with higher cereal density (T+, 3T:1F) showed slightly elevated SIL values. Overall, SIL showed little evidence of systematic plastic adjustment to intercropping conditions.

**Figure 5.**
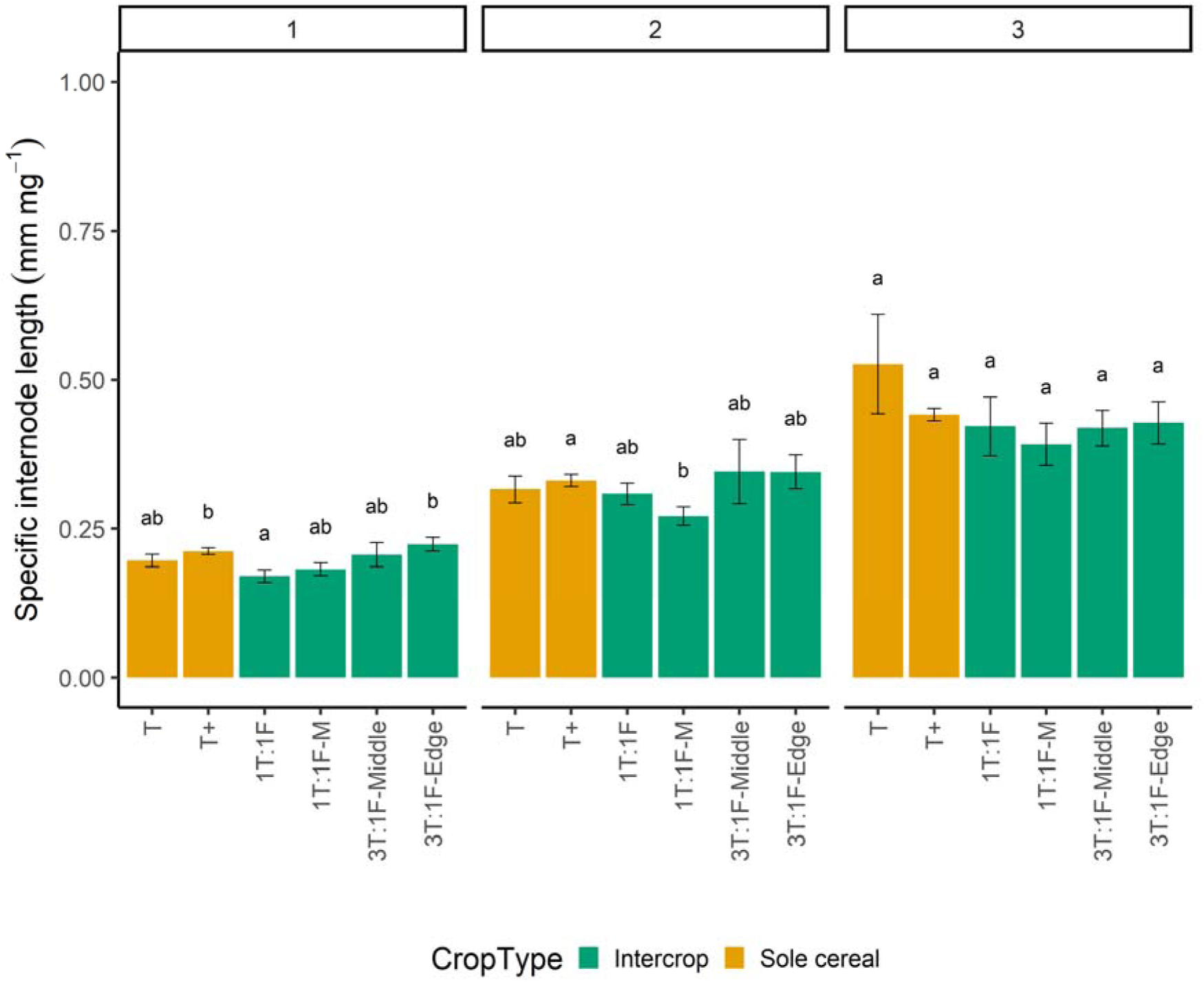
Cereal specific internode length. Specific internode length (SIL; in mm mg^-1^) of cereal plants in sole crops (red) and intercrops (green) across treatments, for the first (1), second (2), and third (3) internode of the plant. Treatments comprised triticale sole crops at standard density (T) and at 1.5 times that density (T+), and intercrops of triticale and faba bean arranged as 1:1 row ratio (1T:1F), mixed within rows (1T:1F-M), and row ratios of 1:3 (1T:3F) and 3:1 (3T:1F). In the 3T:1F treatment, ‘Middle’ and ‘Edge’ indicate the middle row and edge rows of the 3-row strip. Error bars indicate the standard error. Different letters indicate significant differences between treatments at P < 0.05.

### 3.2 Model evaluation

In the second stage of the workflow, we evaluated the FSP model by validating the output against field data and through a sensitivity analysis (Fig. 1b, stage 2). The model reproduced observed growth and morphological patterns well: cereal biomass, grain yield, tiller dynamics, and tiller angles matched field measurements closely in both sole crops and intercrops, though legume sole crop biomass and yield were underestimated and overestimated, respectively. A sensitivity analysis across 72 parameters identified 12 with a sensitivity index above 1.0, of which five were unmeasured, with leaf nitrogen fraction being the most influential for legume biomass. Full evaluation results are provided in Supplement section 4 (Supplementary Figs. S.5-S.10).

### 3.3 Trait derivation

In the third stage of the workflow, we extracted trait values over time of cereal tiller number and angle, SLA, and SIL (Fig. 1a, b, stage 3). To derive cereal trait values across different competitive scenarios, we ran simulations using long-term average radiation and temperature data from 2000-2024, representing typical growing conditions rather than year-specific weather patterns. We simulated six scenarios: a no-competition scenario with isolated plants, a half-density intercrop scenario (weak competition), a weed-infested intercrop scenario (intermediate competition), a weed-infested cereal sole crop scenario (strong competition), and a weed-infested cereal sole crop at double density (extreme competition), while allowing plasticity, to extract trait value temporal dynamics of tiller number and angle, SLA, and SIL (Fig. 6). We ran the model with this long-term average weather data and found that biomass and yield values for sole crops and row intercrops were comparable to the scenario using 2023 radiation and temperature data from the validation step, confirming that using averaged weather conditions produced realistic outputs (Supplementary Figs. S.11 and S.12). Leaf area index (LAI) values showed values increasing for the cereal initially, then dropping as leaves started to senesce (Supplementary Fig. S.13). Legume LAI values increased steadily, as this is a non-determinate species that keeps creating new phytomers with leaves. LAI values were lower in the intercrop, with legume LAI plateauing (Supplementary Fig. S.13b). These values align with values found in literature for triticale (Nielsen et al., 2012) and faba bean (Ebrahimi et al., 2024) at equivalent densities.

**Figure 6.**
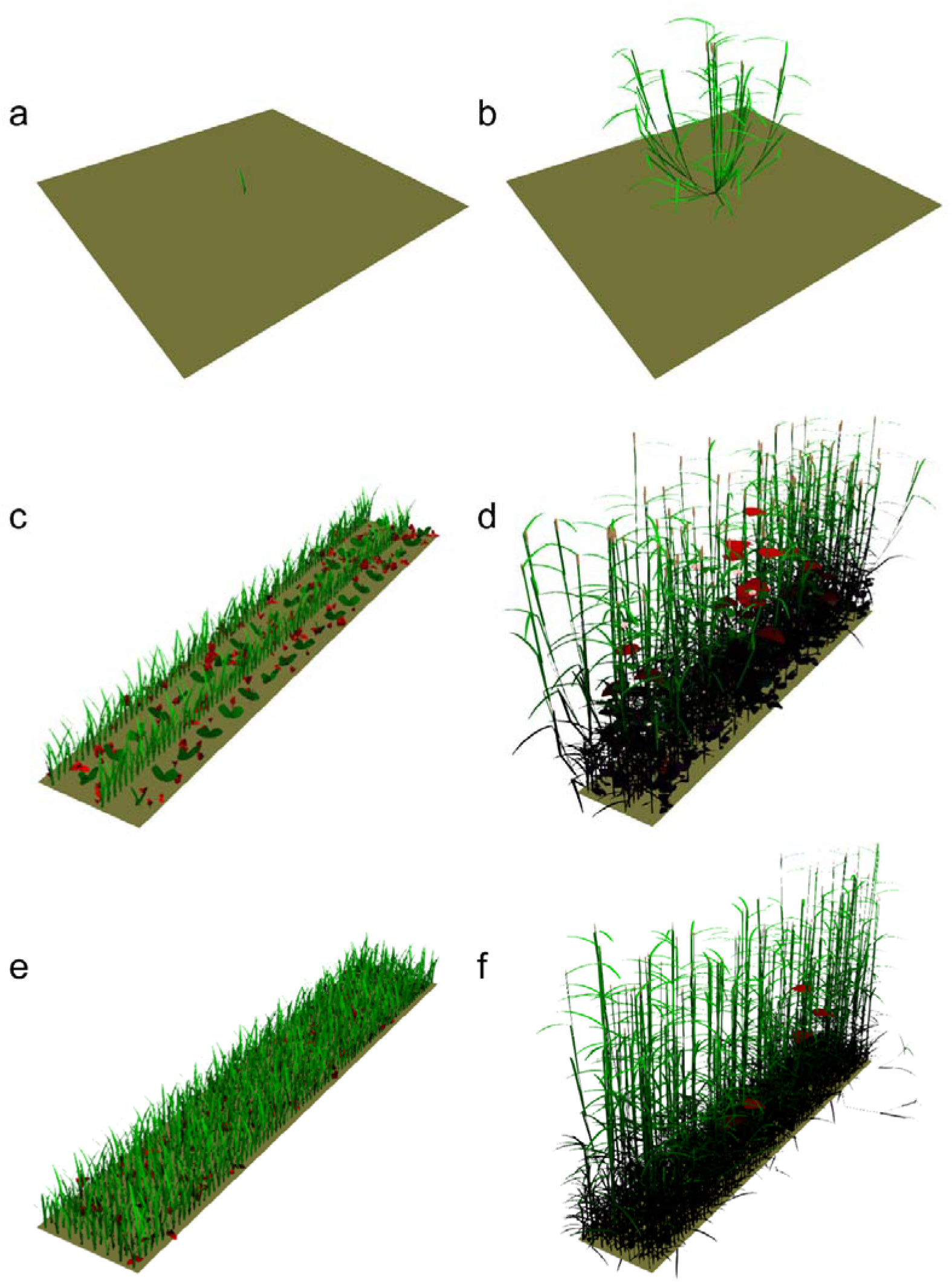
Visual output of several of the crop systems simulated in this study. (a) A solo cereal plant at 25 days after emergence and (b) 85 days after emergence. (c) A cereal-legume weedy intercrop at 25 days after cereal emergence and 17 days after legume emergence; and (d) 85 days after cereal emergence and 77 days after legume emergence. (e) A double-density weedy sole crop cereal at 25 days after emergence, and (f) 85 days after emergence. Weeds are coloured red for increased visibility and emerged simultaneously with cereals. The shade of the colour indicates the proportion of light capture, i.e., a dark or near-black colour indicates low-light capture.

Tiller number and angle showed clear patterns across competitive intensities (Fig. 7a-f, and Supplementary Fig. S.14a-d). As competitive intensity increased from no-competition to extreme-competition, tiller number declined substantially and tillers became more upright with narrower angles relative to the main stem, with intercrop values intermediate between the extremes though more similar to the extreme-competition system. Similarly, SLA and SIL were lowest under no-competition and increased under more competitive conditions, though the extreme-competition and intercrop SIL values were very similar to each other (Fig. 7g-l, and Supplementary Fig. S.14e-h).

**Figure 7.**
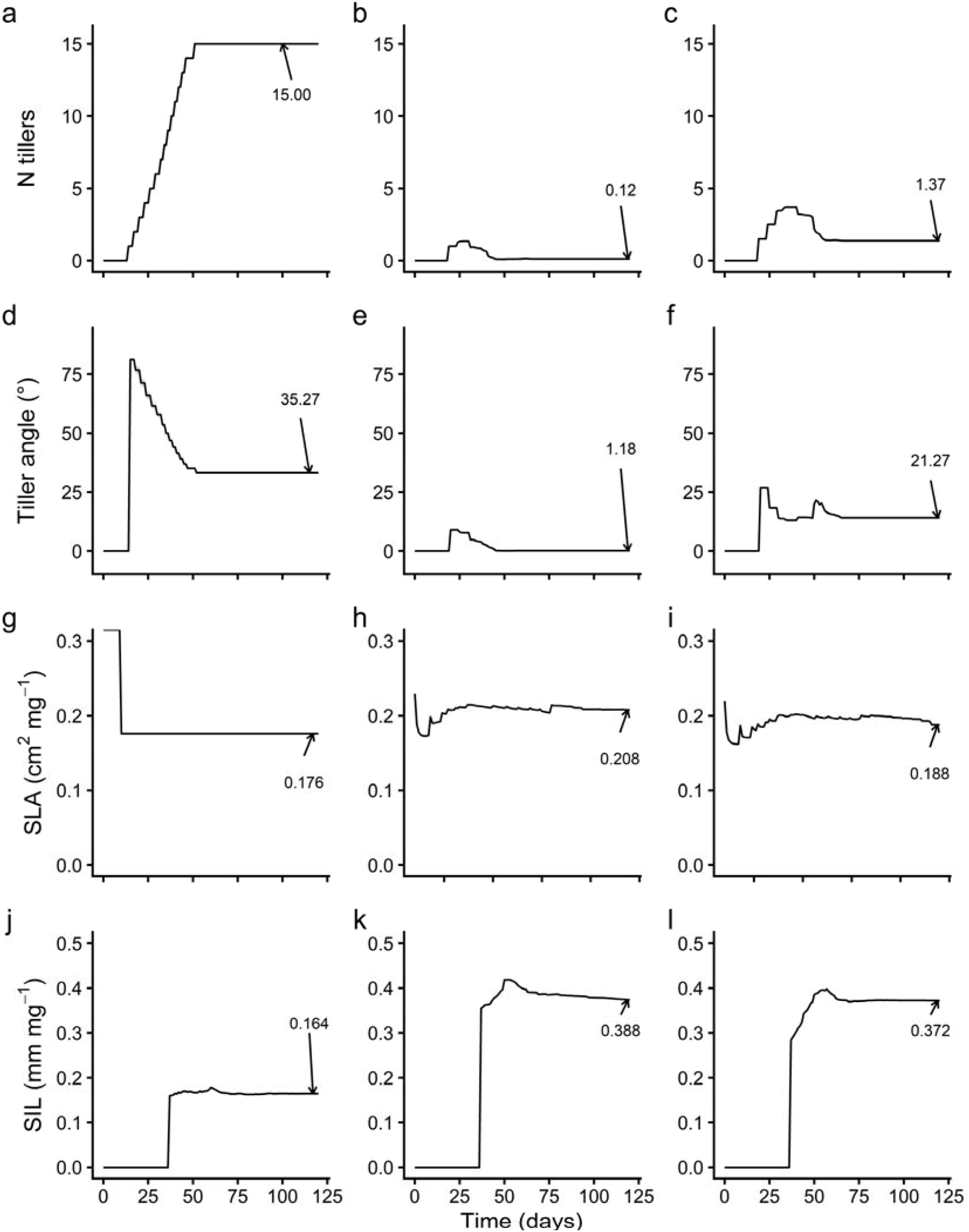
Spatial effect quantification. System outputs for sole crop cereals and intercrops with every combination of sole crop and intercrop cereal phenotypes. (a) Weed biomass (g mL²) over time (days); (b) cereal above-ground biomass (g m^-2^) over time (days); (c) cereal grain biomass at harvest time. Shaded areas and error bars indicate the standard error over three replicate simulations.

### 3.4 Spatial and phenotypic effect separation

In the fourth stage of the workflow, we separated spatial effects (row arrangement, neighbour identity) from phenotypic effects (plastic trait adjustments) by conducting virtual transplant experiments (Fig. 1b, stage 4). We simulated four scenarios: (1) sole crop spatial arrangement with sole-crop trait values (resident), (2) sole crop spatial arrangement with intercrop trait values (transplant), (3) intercrop spatial arrangement with intercrop trait values (resident), and (4) intercrop spatial arrangement with sole-crop trait values (transplant). Comparing resident versus transplant trait values within each spatial arrangement quantifies the benefit of plastic trait adjustment to that competitive environment. The resident trait values consistently outperformed transplanted trait values in both spatial arrangements (Fig. 8). Sole crop systems with sole-crop trait values suppressed weeds most strongly and produced the highest biomass and yield, while intercrop systems with intercrop trait values outperformed those with sole-crop trait values. These patterns aligned with LAI and light absorption dynamics (Supplementary Figs. S.15 and S.16). The consistent advantage of resident over transplanted trait values demonstrates that cereal trait plasticity substantially improved crop biomass, yield, and weed suppression by adjusting trait values to match local competitive conditions.

**Figure 8.**
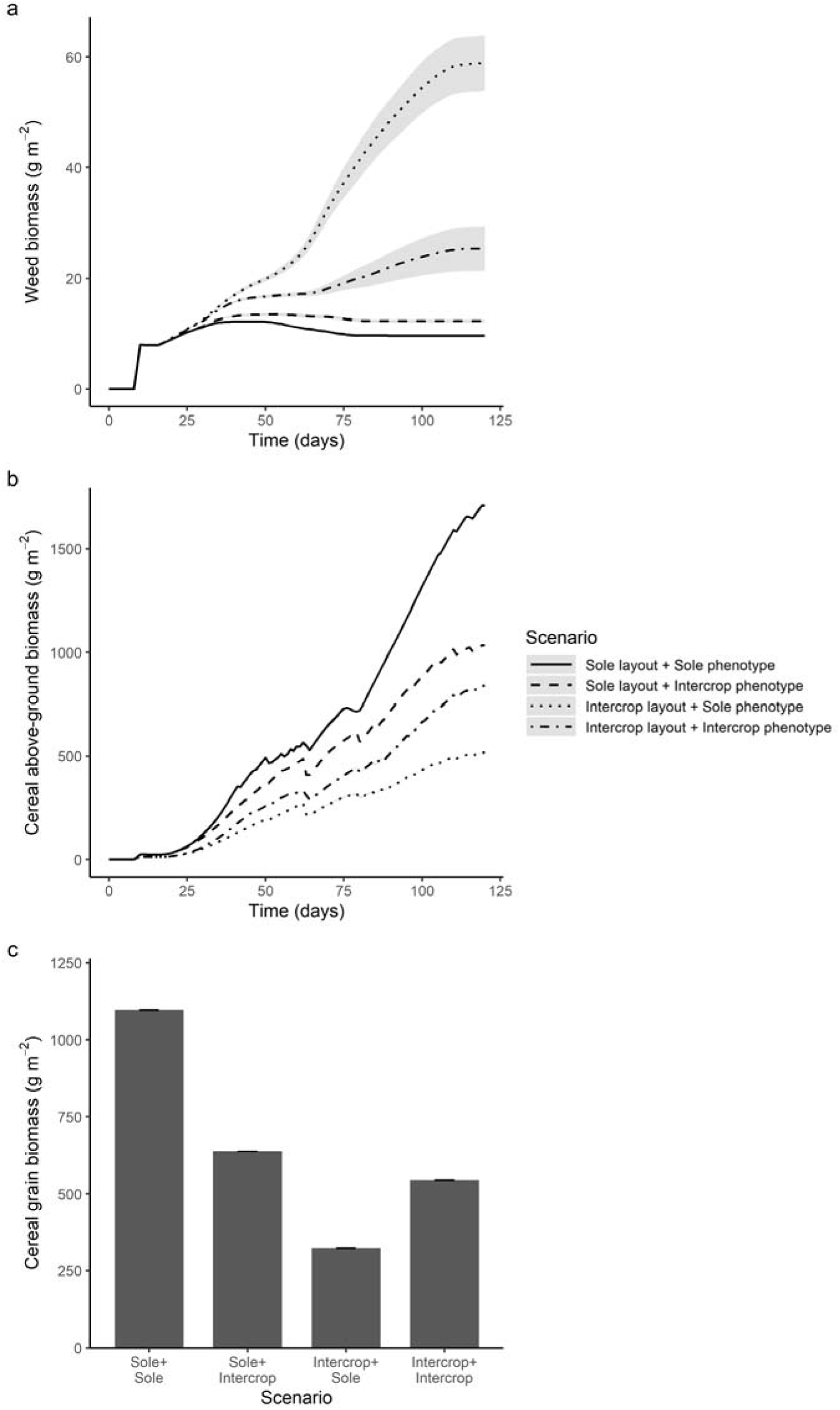
Simulation plastic trait value results. Average trait values over time (days since simulation start) for (a, d, g, e) an isolated cereal plant (i.e., no competition), (b, e, h, k) double-density weed-infested cereal sole crop (i.e., extreme competition), or (c, f, i, l) weed-infested cereal-legume intercrop (i.e., intermediate competition) for (a, b, c) tiller number, (d, e, f) plant mean tiller angle (in degrees), (g, h, i) specific leaf area (SLA; in cm^2^ mg^-1^), and (j, k, l) specific internode length (SIL; in mm mg^-1^). Shaded areas indicate the standard deviation for the three replicate simulations (mostly not visible due to low variation in simulation replicates). The final stable trait values are indicated in the plots.

### 3.5 Trait effect quantification

In the fifth stage of the workflow, we quantified the effects of tiller number, tiller angle, SLA, and SIL on crop productivity and weed suppression in intercrops with and without resource feedbacks (Fig. 1b, stage 5). We tested how individual cereal trait variation affected weed suppression and crop productivity using simulations with predetermined cereal trait trajectories under dynamic (cereal assimilates calculated from light capture) and fixed (cereal assimilates fixed at intercrop levels from stage 3) assimilation modes.

#### 3.5.1 Tiller number

Weed suppression varied with tiller number: extremely high tiller numbers (15 tillers) allowed the most weed biomass, low tiller numbers (0.12 tillers) showed intermediate weed suppression, and moderate tiller numbers (0.57-2.34 tillers) suppressed weeds most effectively (Fig. 9a). Under fixed assimilation, differences between phenotypes diminished overall, but medium-high (2.34 tillers) and medium-low (0.57 tillers) tiller numbers showed slightly lower weed biomass than the intercrop phenotype (1.37 tillers), indicating that the competitive disadvantage of extremely high tiller numbers operates partially through resource-capture feedbacks (Fig. 9b). The slightly enhanced weed suppression of medium-high and medium-low tiller numbers under fixed assimilation suggests two alternative competitive strategies that avoid resource feedback penalties: either moderately high tiller numbers with shorter stems or moderately low tiller numbers with taller stems can provide geometric advantages for weed suppression when resource feedbacks are removed. High-tiller phenotypes produce shorter stems with many tillers, positioning leaves lower and more crowded within the canopy, capturing less light and generating fewer assimilates, which further constrains growth and competitiveness in a reinforcing negative cycle.

**Figure 9.**
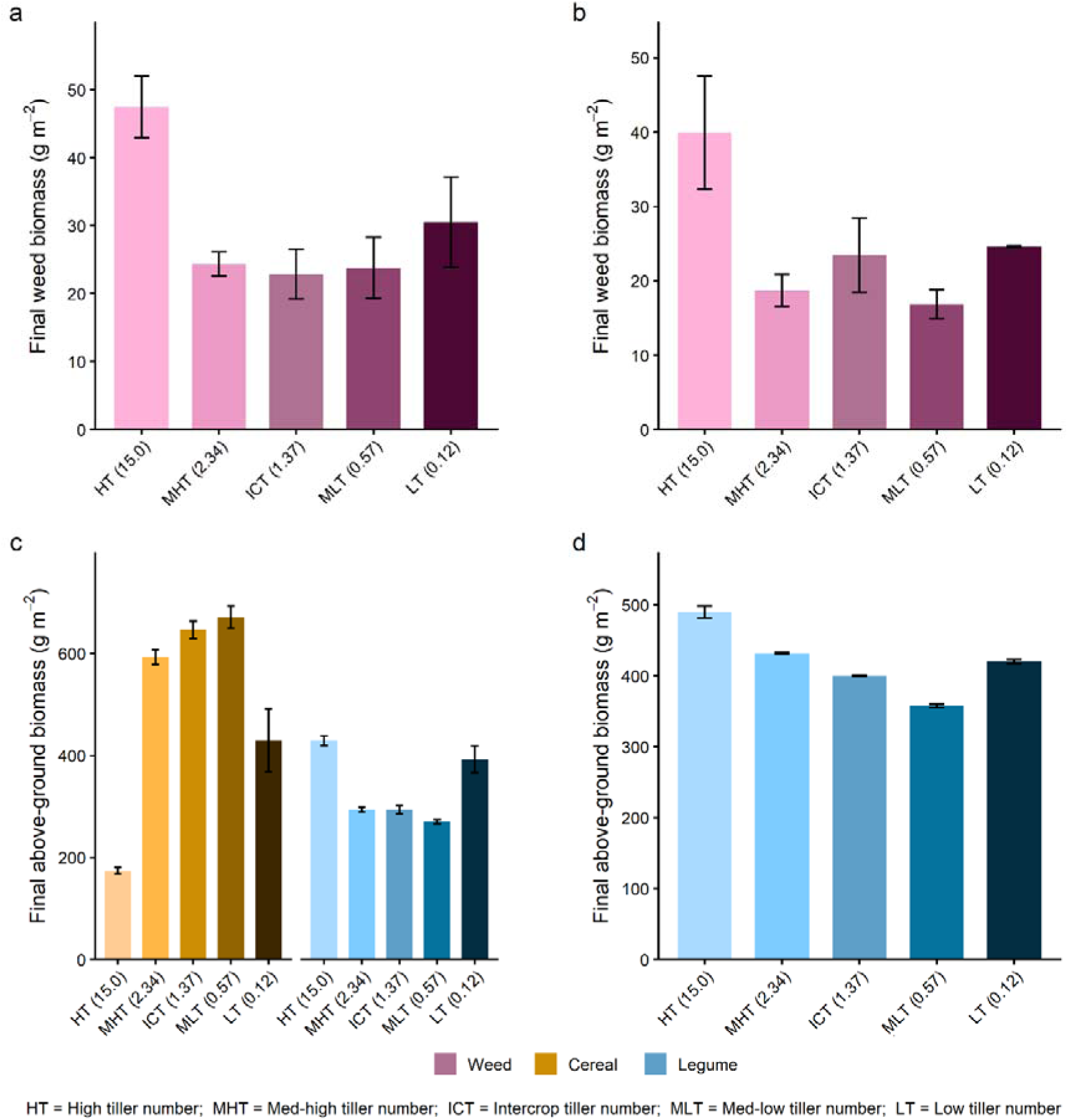
Effect of cereal tillering plasticity on weed suppression and above-ground biomass. Simulations compare two assimilation modes to separate active from passive plasticity: dynamic carbon assimilation calculated from simulated light capture (a, c), representing both active and passive plasticity, and assimilates fixed at intercrop values (b, d), isolating active plasticity effects while removing resource feedbacks. (a, b) Final weed biomass (g mL²) for each tillering phenotype. (c, d) Final above-ground biomass (g mL²) for cereal (orange) and legume (blue). In (d), only legume biomass is shown because cereal biomass does not differ across phenotypes when assimilates are fixed. Tillering phenotypes represent trait trajectories derived from plants grown under different competitive environments: high tillering (HT), medium-high tillering (MHT), intercrop tillering (ICT), medium-low tillering (MLT), and low tillering (LT), with mean final tiller numbers given in parentheses on the x-axis. Colours within each group (pink for weed, orange for cereal, blue for legume) progress from lighter to darker shades from HT to LT. Error bars indicate standard errors.

Biomass patterns began diverging around 65 days after sowing (Supplementary Fig. S.17). Cereal above-ground biomass was substantially lower with the high-tiller phenotype and moderately lower with the low-tiller phenotype, while medium-high, intercrop, and medium-low tiller phenotypes showed similar values with small differences (medium-high < intercrop < medium-low) (Fig. 9c and Supplementary Fig. S.17c). Legume biomass showed the inverse pattern: highest under high-tiller cereal phenotypes and lowest under medium-low tiller phenotypes, with differences diminishing under fixed assimilation (Fig. 9d and Supplementary Fig. S.17d). Grain biomass patterns mirrored above-ground biomass trends (Supplementary Fig. S.17e, f). LAI and absorbed PAR patterns showed that while the high-tiller phenotype maintained higher leaf area than other phenotypes, it absorbed substantially less PAR, indicating inefficient canopy positioning (Supplementary Fig. S.18). This poor light capture efficiency explains both the reduced weed suppression and lower cereal biomass under high tiller numbers: despite producing more leaf area, the high-tiller phenotype captured less light overall due to lower, more crowded leaf positioning.

#### 3.5.2 Tiller angle

For tiller angle, both the intercrop (21.27° from vertical) and wide-angle (35.27°) phenotypes suppressed weeds slightly better than the narrow-angle (1.18°) phenotype under dynamic assimilation, with medium-narrow and medium-wide phenotypes showing intermediate suppression (Fig. 10a). However, under fixed assimilation, this pattern changed, as wide- and narrow-angle phenotypes suppressed weeds slightly more effectively than the intercrop and medium phenotypes, with wide and narrow performing similarly to each other (Fig. 10b). Cereal above-ground biomass, grain yield, and legume biomass and yield showed minimal differences among tiller angle phenotypes (Fig. 10c-f and Supplementary Fig S.19). LAI and absorbed PAR were comparable across all tiller angle phenotypes (Supplementary Fig. S.20), suggesting that competitive differences resulted from leaf positioning within the canopy rather than total leaf area or overall resource availability.

**Figure 10.**
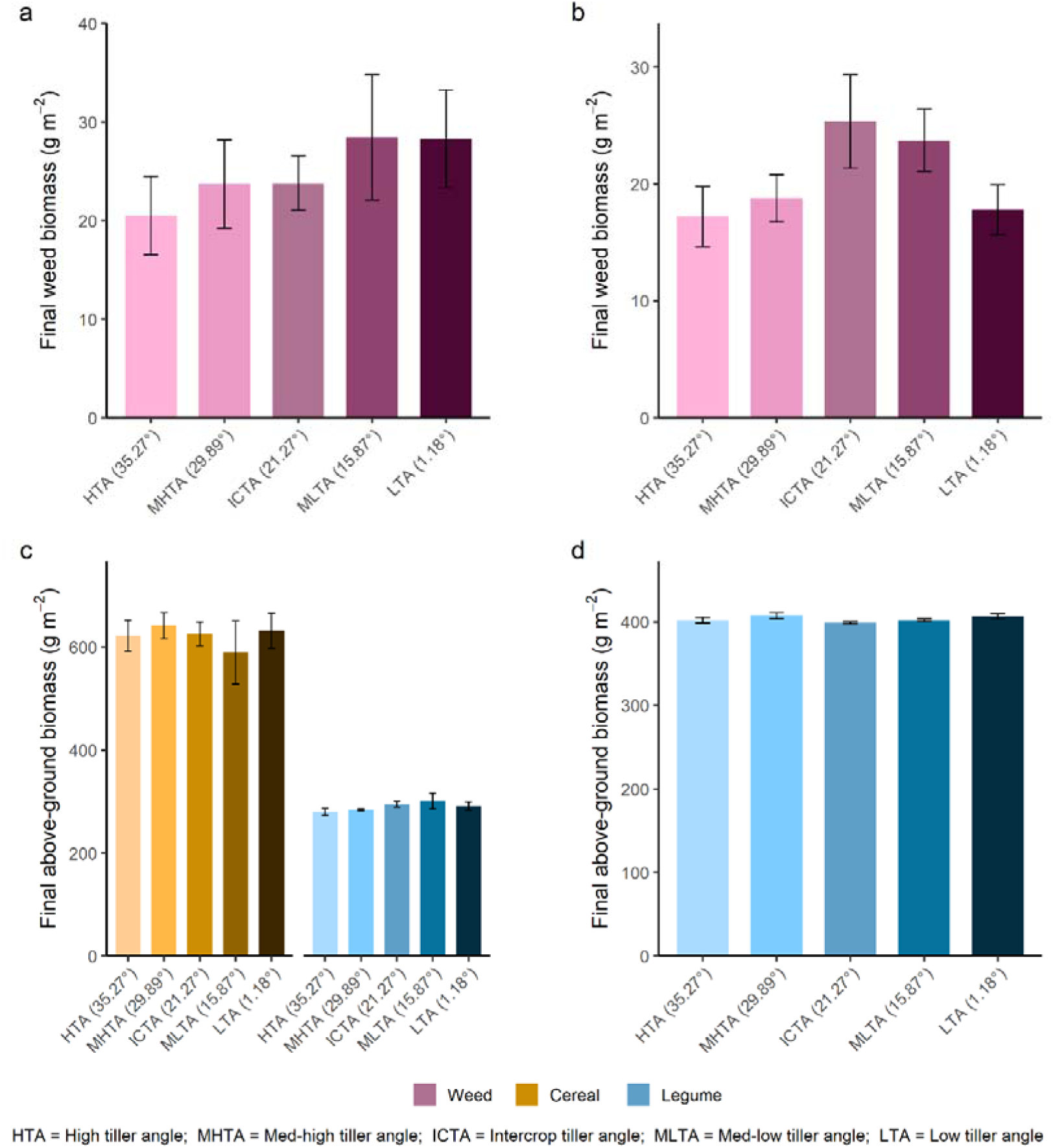
Effect of cereal tiller angle plasticity on weed suppression and above-ground biomass. Simulations compare two assimilation modes to separate active from passive plasticity: dynamic carbon assimilation calculated from simulated light capture (a, c), representing both active and passive plasticity, and assimilates fixed at intercrop values (b, d), isolating active plasticity effects while removing resource feedbacks. (a, b) Final weed biomass (g mL²) for each tiller angle phenotype. (c, d) Final above-ground biomass (g mL²) for cereal (orange) and legume (blue). In (d), only legume biomass is shown because cereal biomass does not differ across phenotypes when assimilates are fixed. Tiller angle phenotypes represent trait trajectories derived from plants grown under different competitive environments: high tiller angle (HTA), medium-high tiller angle (MHTA), intercrop tiller angle (ICTA), medium-low tiller angle (MLTA), and low tiller angle (LTA), with mean final tiller angles given in parentheses on the x-axis. Colours within each group (pink for weed, orange for cereal, blue for legume) progress from lighter to darker shades from HTA to LTA. Error bars indicate standard errors.

#### 3.5.3 Specific leaf area

SLA showed competitive effects that operated primarily through direct leaf area deployment rather than resource-capture feedbacks. Under dynamic assimilation, weed suppression increased progressively with SLA: low-SLA cereals (0.176 cm² mgL¹) allowed the most weed biomass, with medium-low (0.181), intercrop (0.188), and medium-high (0.194) phenotypes showing intermediate suppression, and high-SLA cereals (0.208 cm² mgL¹) suppressing weeds most effectively (Fig. 11a and Supplementary Fig. S.21). Under fixed assimilation, this pattern persisted and the difference between low-SLA cereals and the other SLA phenotypes became more pronounced (Fig. 11b). This indicates that SLA’s competitive advantage operates through direct geometric effects—high-SLA cereals deploy more leaf area per unit mass, physically intercepting more light—and that resource-capture feedback actually partially dampen these differences rather than amplify them. Cereal above-ground biomass and grain yield were slightly lower for the low-SLA phenotype than the other SLA phenotypes (Fig. 11c and Supplementary Fig. S.21c, e). Legume biomass and yield were not substantially affected by cereal SLA phenotype differences (Fig. 11d, f). LAI and PAR absorption patterns showed that the low-SLA phenotype maintained lower leaf area throughout the season with correspondingly lower PAR absorption, benefiting weed and legume light capture (Supplementary Fig. S.22).

**Figure 11.**
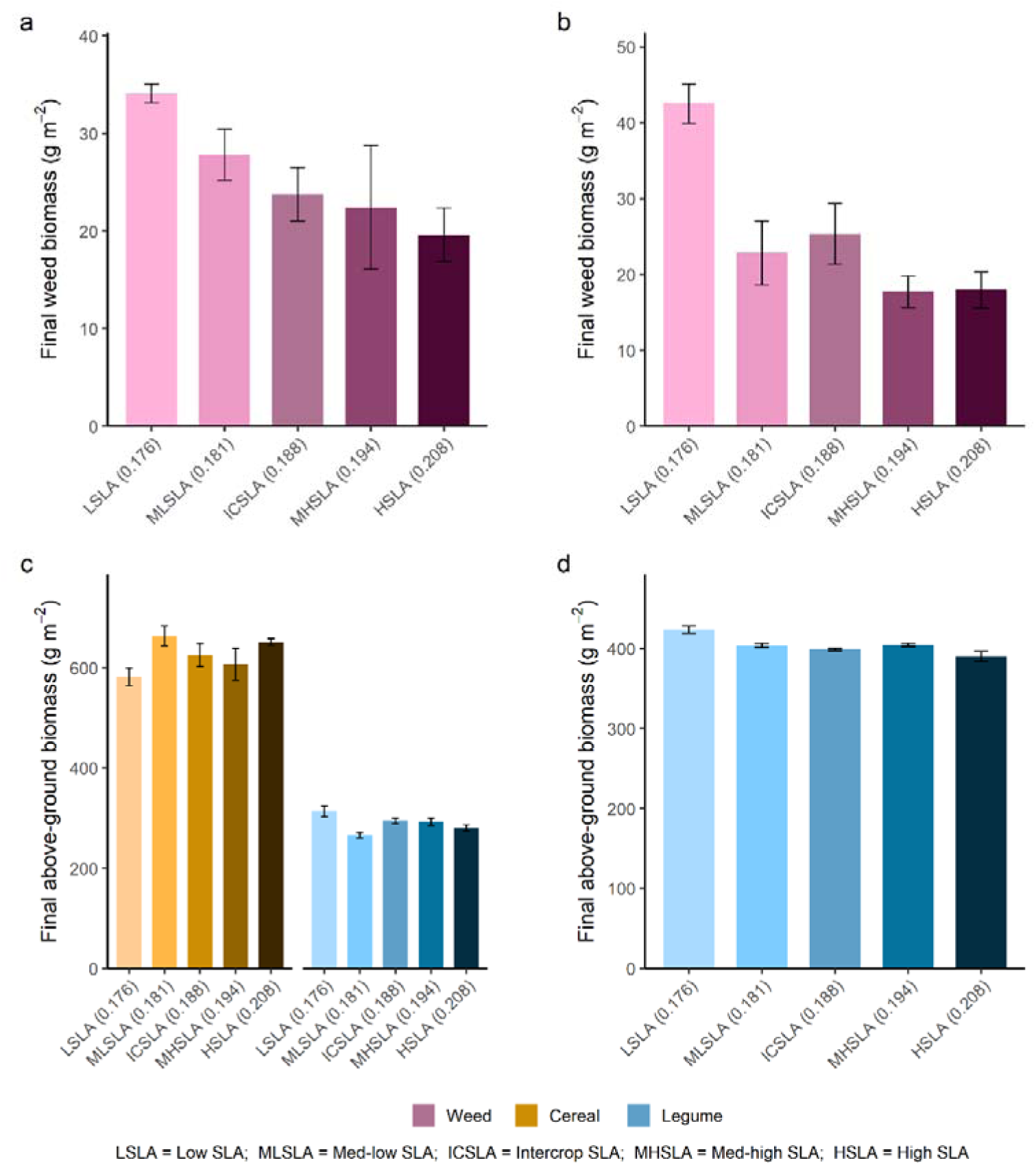
Effect of cereal SLA plasticity on weed suppression and above-ground biomass. Simulations compare two assimilation modes to separate active from passive plasticity: dynamic carbon assimilation calculated from simulated light capture (a, c), representing both active and passive plasticity, and assimilates fixed at intercrop values (b, d), isolating active plasticity effects while removing resource feedbacks. (a, b) Final weed biomass (g mL²) for each SLA phenotype. (c, d) Final above-ground biomass (g mL²) for cereal (orange) and legume (blue). In (d), only legume biomass is shown because cereal biomass does not differ across phenotypes when assimilates are fixed. SLA phenotypes represent trait trajectories derived from plants grown under different competitive environments: high SLA (HSLA), medium-high SLA (MHSLA), intercrop SLA (ICSLA), medium-low SLA (MLSLA), and low SLA (LSLA), with mean final SLA values given in parentheses on the x-axis. Colours within each group (pink for weed, orange for cereal, blue for legume) progress from lighter to darker shades from HSLA to LSLA. Error bars indicate standard errors.

#### 3.5.4 Specific internode length

Under dynamic assimilation, cereal SIL phenotypes had minimal effects on weed biomass, with low-SIL (0.164), medium-low (0.357), intercrop (0.372), medium-high (0.379), and high-SIL (0.388) phenotypes suppressing weeds similarly, and no consistent directional trend across the competition gradient (Fig. 12a and Supplementary Fig. S.23). Under fixed assimilation, differences in weed biomass decreased (Fig. 12b). Cereal above-ground biomass and grain yield were substantially lower for the low-SIL phenotype compared to all other phenotypes under dynamic assimilation, with medium-low SIL showing intermediate values (Fig. 12c and Supplementary Fig,. S.23c, e). Conversely, legume biomass and yield were substantially higher under the low-SIL cereal phenotype, with differences diminishing under fixed assimilation (Fig. 12d, f). Despite the shift in cereal-legume competitive balance under dynamic assimilation, weed biomass remained similar across phenotypes, indicating that legumes had a weed suppressiveness comparable to that of the higher-SIL cereals when not themselves suppressed by competitive cereals. LAI patterns were similar across phenotypes until late in the season, while PAR absorption differed substantially: under dynamic assimilation with the low-SIL phenotype, legumes captured more PAR earlier in the season at the expense of cereal PAR absorption, indicating more competitive leaf positioning by legumes and supporting their higher weed suppressiveness (Supplementary Fig. S.24).

**Figure 12.**
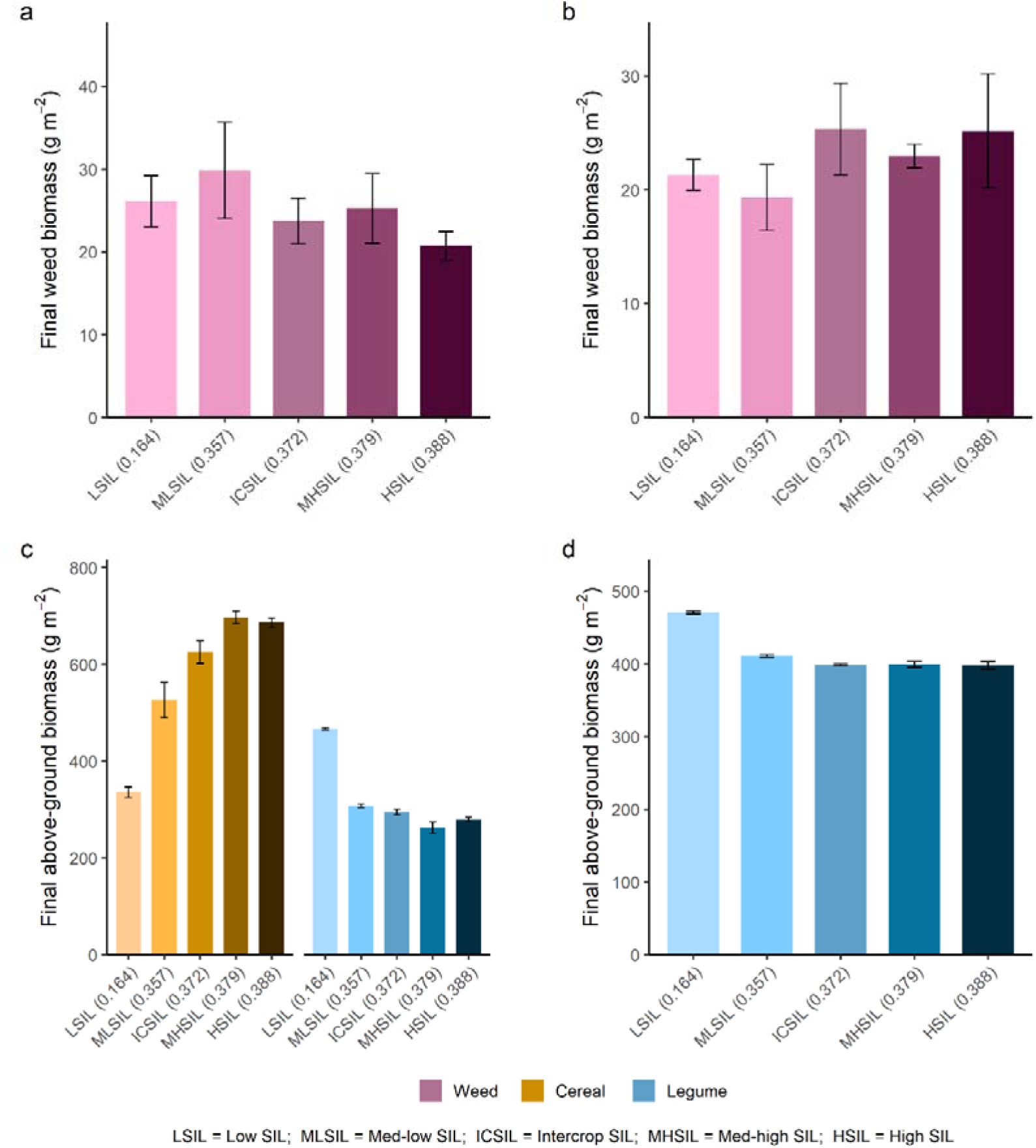
Effect of cereal SIL plasticity on weed suppression and above-ground biomass. Simulations compare two assimilation modes to separate active from passive plasticity: dynamic carbon assimilation calculated from simulated light capture (a, c), representing both active and passive plasticity, and assimilates fixed at intercrop values (b, d), isolating active plasticity effects while removing resource feedbacks. (a, b) Final weed biomass (g mL²) for each SIL phenotype. (c, d) Final above-ground biomass (g mL²) for cereal (orange) and legume (blue). In (d), only legume biomass is shown because cereal biomass does not differ across phenotypes when assimilates are fixed. SIL phenotypes represent trait trajectories derived from plants grown under different competitive environments: high SIL (HSIL), medium-high SIL (MHSIL), intercrop SIL (ICSIL), medium-low SIL (MLSIL), and low SIL (LSIL), with mean final SIL values given in parentheses on the x-axis. Colours within each group (pink for weed, orange for cereal, blue for legume) progress from lighter to darker shades from HSIL to LSIL. Error bars indicate standard errors.

## 4. Discussion

In this study, we combined field experiments with FSP modelling to quantify to what extent cereal traits respond plastically to intercropping environments, and to quantify the extent to and direction in which trait plasticity affects weed suppression and crop productivity. We found that cereal traits differed both in terms of their plastic response patterns and the implications these had on crop competitive performance. For instance, tiller number had an optimum whereby intermediate tiller numbers produced the strongest competitiveness and competitiveness declined towards higher and lower tillering values, consistent with observed field plasticity enabling adjustment to local conditions. SLA showed a progressive pattern where weed suppression increased gradually with SLA value, while SIL showed a saturating pattern where only low values resulted in disproportionate competitive losses, both in agreement with minimal field plasticity. Traits influenced competition through different mechanisms: SLA and SIL operated through direct geometric effects on leaf area deployment and canopy positioning, while tillering affected competition through resource distribution and leaf positioning. Additionally, SIL effects were mediated by cereal-legume competitive interactions where legumes compensated for less competitive cereal phenotypes (those with low SIL) in weed suppression.

### 4.1 Tillering benefits from plasticity due to its optimum competitive response, while SLA and SIL follow progressive and saturating patterns

Field measurements revealed moderate plasticity in tiller number (Fig. 2) and tiller angle (Fig. 3) in response to intercropping. Simulations revealed distinct patterns for these traits. Tiller number showed an optimum at intermediate values: extremely high tiller numbers substantially reduced cereal competitiveness, medium-high and medium-low tiller numbers showed the strongest weed suppression, and very low values showed reduced performance, confirming the role of tillering in balancing resource capture against resource dilution across multiple stems (Fig. 9). Tiller angle showed less clear patterns, with competitive rankings shifting between assimilation modes and relatively small differences in weed suppression overall, making interpretation difficult (Fig. 10).

Tiller number plasticity was not observed in rye and wheat intercrops (Fig. 2), though triticale showed clear plastic responses. Rye was an exceptionally strong weed suppressor across treatments, while wheat performed as a relatively weak competitor (Kottelenberg et al., 2026a). In highly competitive rye systems, intense intraspecific competition may suppress tillering regardless of cropping system. Alternatively, since rye did show increased tiller angles in intercrops, rye’s growth strategy might prioritise optimising tiller positioning over adjusting tiller number. Conversely, wheat plants may experience similar competitive intensity in sole crops and intercrops, providing insufficient environmental contrast to trigger plastic responses in either tiller number or angle, consistent with wheat showing no plasticity in either trait.

Contrasting with the tiller traits, for SLA we did not detect significant differences among field experiment treatments (Fig. 4), while SIL showed only small differences (Fig. 5). In the simulations, SLA showed a progressive pattern where weed suppression increased gradually with SLA value, while SIL showed a saturating pattern where only low values resulted in disproportionate competitive losses (Figs. 11 and 12). These effects operated through direct geometric mechanisms—insufficient leaf area deployment (low SLA) or inadequate canopy height (low SIL)—rather than through resource-capture feedbacks, as evidenced by the persistence or amplification of competitive differences when resource feedbacks were eliminated through fixed resource assimilation. Interestingly, cereal phenotypes with low SIL values were outcompeted by legumes, which subsequently compensated for the cereals in terms of weed suppression. This aligns with field observations that, although the cereal canopy is dominant early in the season, legumes have the potential to become dominant later in the season (Kottelenberg et al., 2026b), with transition timing possibly partially determined by cereal height.

These distinct patterns—optimisation for tillering, a progressive gradient for SLA, and a saturating pattern for SIL—may explain why in the field we observed stronger plasticity in tiller traits than in either SLA or SIL. Tillering requires precise adjustment to local competitive conditions to avoid penalties from excessive or insufficient tiller numbers, making plasticity advantageous. By contrast, SIL primarily requires staying above a critical minimum, while any increase in SLA provides some competitive benefit, reducing selective pressure for precise plastic adjustment in either trait. The minimal SIL plasticity is somewhat surprising given the substantial internode elongation plasticity documented in many plant species (Gautrat et al., 2025; Schmitt et al., 2003; Morelli and Ruberti, 2000; Schmitt and Wulff, 1993; Smith, 1982), but in cereals, competitive canopy height is determined by multiple traits including leaf length and tillering, potentially reducing the relative importance of SIL plasticity. Trade-offs may further constrain plasticity: excessive stem elongation increases lodging risk (Sparkes and King, 2008) while high-SLA leaves suffer increased herbivore damage (Caldwell et al., 2016), favouring moderate adjustments over dramatic phenotypic shifts (Schneider et al., 2022; Niinemets, 2020).

The absence of observed SLA and SIL plasticity may additionally reflect modern cereal breeding for monoculture conditions, which has minimised plastic responses that would cause excessive intraspecific competition in high-density systems (Vandermeer and Perfecto, 2025; Bourke et al., 2021; Anten and Vermeulen, 2016), or detection limitations inherent to field conditions. Greenhouse studies consistently detect SLA and SIL plasticity under controlled conditions (Golan et al., 2023; Arenas-Corraliza et al., 2021; Dufour et al., 2012), while field experiments often find minimal differences (Panozzo et al., 2025; Dos Santos et al., 2023; Lafarge and Hammer, 2002). However, even if small plastic adjustments do occur in the field, our simulations suggest their competitive benefits would be modest: for SIL, only values below a critical threshold substantially affect competitiveness, while for SLA, the progressive response means small adjustments yield only small gains.

### 4.2 Tiller number shows resource-capture feedback effects while SLA and SIL operate through geometric mechanisms

To investigate whether trait effects on competition operate through resource-capture feedbacks or through direct geometric positioning, we compared simulations under dynamic assimilation (default setting, where light capture determines available assimilates), where trait variations affect both architecture and resource availability, versus fixed assimilation (daily available assimilates are given as input and fixed to the intercrop-phenotype values, and are not affected by light capture), where total cereal assimilates per plant were held constant, isolating geometric effects. Under dynamic assimilation, changes in cereal plant architecture influence light capture and thereby alter available resources for growth in reinforcing feedback cycles. Under fixed assimilation, these whole-plant resource feedbacks were eliminated, though within-plant source-sink dynamics among individual organs continued to operate.

Under dynamic assimilation, extremely high tiller numbers (15 tillers) substantially reduced cereal competitiveness compared to intermediate values (Fig. 9a). Under fixed assimilation, this disadvantage intensified (Fig. 9b), indicating that resource feedbacks partially compensated for high tiller numbers under dynamic conditions. With dynamic assimilation, high-tiller plants could offset their poor leaf positioning in the light gradient by capturing additional light through their numerous tillers. In our simulations this compensation was insufficient to overcome the fundamental positioning problem. When assimilates were fixed regardless of light capture, this compensatory mechanism disappeared, revealing the full extent of the geometric disadvantage. These patterns are consistent with findings that architectural differences can create competitive effects through feedback loops (Bongers et al., 2017; Anten et al., 2005), in this case demonstrating how feedbacks can partially mask underlying geometric constraints.

In contrast, SLA showed the opposite pattern. Under dynamic assimilation, low-SLA cereals allowed substantially more weed biomass than high-SLA and intercrop phenotypes, and under fixed assimilation, this difference became even more pronounced (Fig. 11). This indicates that SLA operates primarily through direct geometric effects—insufficient leaf area deployment—with resource feedbacks actually dampening rather than amplifying competitive differences. Similarly, SIL showed patterns largely independent of resource feedbacks (Fig. 12). Tiller angle showed less clear patterns with competitive rankings shifting between assimilation modes (Fig. 10), making interpretation difficult. The plastic tiller angle response we observed in field experiments (Fig. 3) may represent adjustment to local competitive conditions, consistent with studies showing tiller angle plasticity improves grain yield under varying densities (Montazeaud et al., 2020), though the specific mechanisms remain unclear from our simulations.

### 4.3 Plant spatial arrangement determines which traits matter most for competition

Our findings partially align with studies of strip intercropping systems where tiller number had substantial competitive impacts (Zhu et al., 2015), confirming that tillering plays an important role across different spatial arrangements. However, the way tillering affects light capture competition may differ between systems. In cropping systems with wide strips, strong environmental gradients develop at strip boundaries, creating distinct edge and inner zones where tillering plasticity becomes particularly valuable: edge-row cereal plants can expand laterally into available space through increased tiller production and wider tiller angles, while plants in the middle of a strip will experience much more intraspecific competition and will subsequently produce fewer tillers at potentially narrower angles. In our row intercropping system, where species alternate every 12.5 cm, such large-scale spatial gradients and edge opportunities are absent. Instead, every cereal plant experiences similar competitive environments from neighbouring rows. This makes avoiding excessive tiller numbers more important than exploiting spatial heterogeneity through lateral expansion, suggesting that a high-tillering phenotype might be more suited for edge rows of strip intercrops. Barillot et al. (2019) similarly demonstrated through FSP modelling that the competitive advantage of vertical versus horizontal leaf orientations in wheat shifted with plant density and mixture composition, with more horizontal architectures becoming more competitive in mixed stands than in pure stands. The patterns we observed for SLA and SIL—a progressive response for SLA and a saturating response for SIL, both requiring adequate values to maintain competitive ability—likely apply across spatial arrangements including strip systems. Regardless of row versus strip configuration, insufficient leaf area deployment or inadequate plant height would impair competitive ability, depending on the companion crop. However, the absence of strong horizontal gradients in row intercrops may reduce both the opportunity for plastic SLA and SIL adjustment and the competitive advantage such adjustments might provide. In strip systems with distinct edge zones, plants might encounter sufficiently different light environments to trigger plastic responses.

This context-dependence has important practical implications: there is unlikely to be a universal “intercrop ideotype” because the relative importance of traits depends on spatial design. This supports the view that spatial arrangement determines the relevance of trait plasticity, which matters more for competitive outcomes in spatially segregated systems than in intensely mixed arrangements (Bongers et al., 2025).

### 4.4 Model limitations and future research needs

Our simulations revealed distinct mechanisms through which cereal traits influence competitive outcomes, but several model limitations require acknowledgment. Most fundamentally, we cannot be certain the model accurately represents field dynamics, particularly in complex intercrop systems where multiple species interact. Our representation of weeds was necessarily simplified: although individual weed plants varied stochastically in leaf and stem characteristics, they do not approach the diversity of real weed communities where different species with distinct competitive strategies emerge at different times throughout the season (Barroso et al., 2015; Gunton et al., 2011). Early-emerging weeds compete primarily during cereal-dominated phases, while late-emerging weeds may encounter more legume competition, potentially shifting which cereal traits matter most.

Additionally, our stage 4 and 5 simulations imposed uniform cereal phenotypes for the focal cereal traits to isolate individual trait effects, eliminating within-population variability that characterises real crop stands. Natural phenotypic variation could create competitive hierarchies where a few dominant individuals disproportionately suppress weeds (Huel and Hucl, 1996). This potentially changes how population-mean trait values translate to system performance. Fixing traits at the individual plant level (maintaining realistic population variation) rather than at the population level (where all plants share identical values) would enable more accurate quantification of plasticity effects. Alternatively, or complementarily, field experiments representing a range of competitive pressures could validate whether the patterns we observed in simulations—optimisation dynamics for tillering, progressive and saturation patterns for SLA and SIL—manifest under realistic conditions with natural phenotypic variation and diverse weed communities.

The mechanisms we used to model plasticity represent simplified approximations of complex biological processes and we may thus have oversimplified some responses to competitive signals. To more realistically quantify plasticity contributions, future work should better parameterise and calibrate the plastic responses observed in the field—particularly tiller number and angle plasticity—ensuring modelled mechanisms match field observations. Additionally, our model captures only above-ground light competition, excluding belowground interactions for water and nutrients that might strongly affect competitive outcomes (Kaur et al., 2018). Whether the patterns we observed—particularly SLA feedback dependence and SIL saturating effects—extend to soil resource-limited environments where below-ground competition dominates remains unknown. Future model development incorporating root trait variation, water and nutrient competition, and more sophisticated plasticity mechanisms would determine whether our findings generalise beyond light-limited conditions.

### 4.5 Implications for intercrop breeding and design

Given model uncertainties, both breeding and system design offer complementary pathways toward improved intercrop performance, each with distinct advantages and limitations. Breeding for intercropping presents inherent complexity because it requires optimising performance in mixed stands rather than maximising individual crop productivity (Bourke et al., 2021). Our findings add to this complexity: plasticity can have both positive effects (optimising cereal performance through tillering adjustment) and create trade-offs (highly competitive cereals suppressing legumes versus legume compensation maintaining weed control when cereals are less dominant). Trait effects operate through multiple mechanisms—resource-capture feedbacks and optimisation for tillering, geometric effects for SLA, cereal-legume competitive dynamics for SIL—meaning plasticity effects depend on context in ways that complicate breeding for intercropping (Bongers et al., 2025). Evidence that tillering mainly affects performance differences between sole crops and intercrops (Zhu et al., 2015), together with our results suggesting an optimisation mechanism, indicate that breeding for greater tillering plasticity would improve adaptation of tiller number to local competitive conditions across crop systems and environments.

System design adjustments—modifying species composition, species proportions, row distance, or sowing densities—offer approaches that work with existing cultivar diversity without requiring genetic modification, and can be implemented and refined immediately based on local performance (Kottelenberg et al., 2026a). This makes design particularly valuable given breeding target uncertainties: if predictions about optimal trait values prove inaccurate, design can adapt quickly, whereas breeding commitments are long-term. However, design cannot fully overcome fundamental limitations in cultivar trait profiles, and both approaches are ultimately subject to genotype × environment × management (G×E×M) constraints. Breeding for appropriate trait plasticity could therefore complement design: if plastic responses preserve performance across environments and cropping systems, they can increase robustness. Combining both—using design to identify which trait profiles perform best under different spatial arrangements, then targeting those profiles in breeding programmes—offers synergistic benefits. Field experiments crossing specific design choices with cultivars differing in trait values and plasticity, conducted across diverse environments, would quantify trade-offs between weed suppression and crop yields and inform both strategies directly. Given the large number of treatment combinations involved, FSP modelling could guide experimental design and prioritise treatment selection, though model refinement and validation should precede any firm conclusions for breeding or design optimisation.

This work represents progress toward mechanistic understanding of trait effects in intercrop systems: it demonstrates how FSP modelling can investigate plasticity in cereals and quantify trait contributions to weed suppression and crop performance. However, the specific patterns we observed require validation through field experiments before guiding breeding or design decisions. Integrating FSP modelling with multi-location field experiments could identify design principles that work across soil types, climates, and management regimes, or conclude that different environments require different models altogether.

## Supplementary data

The following supplementary data are available:

Table S.1: Seeding rates and densities of the experimental plots.

Table S.2: Cereal parameters for the functional-structural plant model.

Table S.3: Legume parameters for the functional-structural plant model.

Table S.4: Weed parameters for the functional-structural plant model.

Figure S.1: Average daily temperature and daily total precipitation of experimental years.

Figure S.2: Experimental setup of the 2022 experiment.

Figure S.3: Experimental setup of the 2023 experiments.

Figure S.4: Experimental setup of the 2024 experiment.

Figure S.5: Sensitivity analysis results of the functional-structural plant model.

Figure S.6: Biomass validation of the functional-structural plant model.

Figure S.7: Yield validation of the functional-structural plant model.

Figure S.8: Tiller number validation of the functional-structural plant model.

Figure S.9: Tiller angle validation of the functional-structural plant model.

Figure S.10: Functional-structural plant model plant height distribution.

Figure S.11: Biomass validation of the functional-structural plant model with average temperature input.

Figure S.12: Yield validation of the functional-structural plant model with average temperature input.

Figure S.13: Leaf area index validation of the functional-structural plant model.

Figure S.14: Simulation plastic trait value results of strong and weak competitive environments.

Figure S.15: Spatial effect leaf area index comparison.

Figure S.16: Spatial effect photosynthetically active radiation comparison.

Figure S.17: Effect of cereal tillering plasticity on weed suppression and above-ground biomass.

Figure S.18: Effect of tillering plasticity on leaf area and light absorption.

Figure S.19: Effect of cereal tiller angle plasticity on weed suppression and above-ground biomass.

Figure S.20: Effect of tiller angle plasticity on leaf area and light absorption.

Figure S.21: Effect of cereal specific leaf area plasticity on weed suppression and above-ground biomass.

Figure S.22: Effect of specific leaf area plasticity on leaf area and light absorption.

Figure S.23: Effect of cereal specific internode length plasticity on weed suppression and above-ground biomass.

Figure S.24: Effect of specific internode length plasticity on leaf area and light absorption.

## Acknowledgements

We would like to thank Peter van der Putten and Eva Goudsmit for their knowledge and skills in the planning and execution of the field experiments conducted in this research. We would like to thank Flores Eversdijk, Pim Vrehen, Kshipra Shipra, Nagarjuna Raavi, Lisa Kiel, and Maylis Levallet for their work done in the field experiments. Figure 1 was created in BioRender, Kottelenberg, D. (2026) https://BioRender.com/dlrt7x7.

## Author contribution

DBK: conceptualisation; DBK, AM: methodology; DBK, AM: software; DBK: validation; DBK: formal analysis; DBK: investigation; AM: resources; DBK: writing – original draft; DBK, AM, NPRA, LB, JBE: writing – review and editing; DBK: visualisation; NPRA, LB, JBE: supervision.

## Data availability

The datasets generated and/or analysed during the current study along with the code used to analyse the data are available in a public GitHub, github.com/David-BK/Cereal-trait-plasticity-and-weed-suppression-in-cereal-legume-intercrops---data-and-analysis, and through a public dataset at doi: 10.17026/LS/1N9E5E.

## Footnotes

1 git.wur.nl/jochemevers1/FSPM_BASIC

## References

Ajal J, Kiaer LP, Pakeman RJ, Scherber C, Weih M. 2022. Intercropping drives plant phenotypic plasticity and changes in functional trait space. Basic and Applied Ecology doi: 10.1016/j.baae.2022.03.009

Arasan AP, Radhamani S, Sudha M, Sundarrajan RV, Prasad SA. 2025. Development of herbicide resistant weeds and their impact on future weed management. Discover Applied Sciences doi: 10.1007/s42452-025-07680-0

Anten NPR, Casado-Garcia R, Nagashima H. 2005. Effects of mechanical stress and plant density on mechanical characteristics, growth, and lifetime reproduction of tobacco plants. The American Naturalist doi: 10.1086/497442

Anten NPR, Vermeulen PJ. 2016. Tragedies and crops: understanding natural selection to improve cropping systems. Trends in Ecology and Evolution doi: 10.1016/j.tree.2016.02.010

Arenas-Corraliza MG, Rolo V, Lopez-Diaz ML, Moreno G. 2021. Wheat and barley can increase grain yield in shade through acclimation of physiological and morphological traits in Mediterranean conditions. Scientific Reports doi: 10.1038/s41598-019-46027-9

Barillot R, Escobar-Gutierrez AJ, Fournier C, Huynh P, Combes D. 2014. Assessing the effects of architectural variations on light partitioning within virtual wheat-pea mixtures. Annals of Botany doi: 10.1093/aob/mcu099

Barillot R, Chambon C, Fournier C, Combes D, Pradal C, Andrieu B. 2019. Investigation of complex canopies with a functional-structural plant model as exemplified by leaf inclination effect on the functioning of pure and mixed stands of wheat during grain filling. Annals of Botany doi: 10.1093/aob/mcy208

Barroso J, Miller ZJ, Lehnhoff EA, Hatfield PG, Menalled FD. 2015. Impacts of cropping system and management practices on the assembly of weed communities. Weed Research doi: 10.1111/wre.12155

Baumann DT, Kropff MJ, Bastiaans L. 2000. Intercropping leeks to suppress weeds. Weed Research 40, 359–374

Bedoussac L, Journet E, Hauggaard-Nielsen H, Naudin C, Corre-Hellou G, Prieur L, Jensen E, Justes E. 2014. Eco-functional intensification by cereal-grain legume intercropping in organic farming systems for increased yields, reduced weeds and improved grain protein concentration. In: Bellon S, Penvern S, eds. Organic Farming, Prototype for Sustainable Agricultures. Dordrecht: Springer, 47–63.

Bongers FJ, Pierik R, Anten NPR, Evers JB. 2017. Subtle variation in shade avoidance responses may have profound consequences for plant competitiveness. Annals of Botany doi: 10.1093/aob/mcx151

Bongers FJ, Evers JB, Anten NPR. 2025. Plastic responses for intercrop functioning. Sustainable Agriculture doi: 10.1038/s44264-025-00048-2

Bourke PM, Evers JB, Bijma P, Van Apeldoorn DF, Smulders MJM, Kuyper TW, Mommer L, Bonnema G. 2021. Breeding beyond monoculture: putting the ”intercrop” into crops. Frontiers in Plant Science doi: 10.3389/fpls.2021.734167

Box GEP, Cox DR. 1964. An analysis of transformations. Journal of the Royal Statistical Society. Series B (Methodological) 26, 211–252.

Brooker RW, Bennett AE, Cong WF, et al. 2015. Improving intercropping: a synthesis of research in agronomy, plant physiology and ecology. New Phytologist doi: 10.1111/nph.13132

Bybee-Finley KA, Ryan MR. 2018. Advancing intercropping research and practices in industrialized agricultural landscapes. Agriculture doi: 10.3390/agriculture8060080

Caldwell E, Read J, Sanson GD. 2016. Which leaf mechanical traits correlate with insect herbivory among feeding guilds? Annals of Botany doi: 10.1093/aob/mcv178

Cao Y, Zhong Z, Wang H, Shen R. 2022. Leaf angle: a target of genetic improvement in cereal crops tailored for high-density planting. Plant Biotechnology Journal doi: 10.1111/pbi.13780

Dos Santos Neto A, Panozzo A, Piotto S, Mezzalira G, Furlan L, Vamerali T. 2023. Screening old and modern wheat varieties for shading tolerance within a specialized poplar plantation for agroforestry farming systems implementation. Agroforestry Systems doi: 10.1007/s10457-024-00956-1

Dufour L, Metay A, Talbot G, Dupraz C. 2012. Assessing light competition for cereal production in temperate agroforestry systems using experimentation and crop modelling. Journal of Agronomy and Crop Science doi: 10.1111/jac.12008

Ebrahimi N, Salihy AR, Alipour S, Mozafari SH, Aliyar J, Darwish I. 2024. Prediction of distribution of dry matter and leaf area of faba bean (Vicia faba) using nonlinear regression models. Agricultural Research doi: 10.1007/s40003-024-00700-2

Evers JB, Vos J, Andrieu B, Struik PC. 2006. Cessation of tillering in spring wheat in relation to light interception and red : far-red ratio. Annals of Botany doi: 10.1093/aob/mcl020

Evers JB, Vos J, Fournier C, Andrieu B, Chelle M, Struik PC. 2007. An architectural model of spring wheat: evaluation of the effects of population density and shading on model parameterization and performance. Ecological Modelling doi: 10.1016/j.ecolmodel.2006.07.042

Evers JB, Vos J, Yin X, Romero P, van der Putten PEL, Struik PC. 2010. Simulation of wheat growth and development based on organ-level photosynthesis and assimilate allocation. Journal of Experimental Botany doi: 10.1093/jxb/erq025

Evers JB, Van der Krol AR, Vos J, Struik PC. 2011. Understanding shoot branching by modelling form and function. Trends in Plant Science doi: 10.1016/j.tplants.2011.05.004

Evers JB, Van der Werf W, Stomph TJ, Bastiaans L, Anten NPR. 2019. Understanding and optimizing species mixtures using functional-structural plant modelling. Journal of Experimental Botany doi: 10.1093/jxb/ery288

Franklin KA. 2008. Shade avoidance. New Phytologist doi: 10.1111/j.1469-8137.2008.02507.x

Gautrat P, Matton SEA, Oskam L, Shetty SS, Van der Velde KJ, Pierik R. 2025. Lights, location, action: shade avoidance signalling over spatial scales. Journal of Experimental Botany doi: 10.1093/jxb/erae217

Golan G, Abbai R, Schnurbusch T. 2023. Exploring the trade-off between individual fitness and community performance of wheat crops using simulated canopy shade. Plant, Cell and Environment doi: 10.1111/pce.14499

Gu C, Bastiaans L, Anten NPR, Makowski D, van der Werf W. 2021. Annual intercropping suppresses weeds: a meta-analysis. Agriculture, Ecosystems and Environment doi: 10.1016/j.agee.2021.107658

Gu C, van der Werf W, Bastiaans L. 2022. A predictive model for weed biomass in annual intercropping. Field Crops Research doi: 10.1016/j.fcr.2021.108388

Gunton RM, Petit S, Gaba S. 2011. Functional traits relating arable weed communities to crop characteristics. Journal of Vegetation Science 22, 541–550.

Hauggaard-Nielsen H, Jornsgaard B, Kinane J, Jensen ES. 2008. Grain legume-cereal intercropping: the practical application of diversity, competition and facilitation in arable and organic cropping systems. Renewable Agriculture and Food Systems doi:

10.1017/S1742170507002025

Huel DG, Hucl P. 1996. Genotypic variation for competitive ability in spring wheat. Plant Breeding doi: 10.1111/j.1439-0523.1996.tb00927.x

Kahlen K, Wiechers D, Stutzel H. 2008. Modelling leaf phototropism in a cucumber canopy. Functional Plant Biology doi: 10.1071/FP08034

Kaur S, Kaur R, Chauhan BS. 2018. Understanding crop-weed-fertilizer-water interactions and their implications for weed management in agricultural systems. Crop Protection doi: 10.1016/j.cropro.2017.09.011

KNMI. 2025. Daggegevens van het weer in Nederland. Accessed July 2025.

Kottelenberg DB, Evers JB, Anten NPR, Rangs M, Bastiaans L. 2025. Effects of cereal-legume intercrop system design on weed suppression. Research Square doi: 10.21203/rs.3.rs-5455247/v3. [Preprint].

Kottelenberg DB, Evers JB, Anten NPR, Bastiaans L. 2026a. Managing species dominance in cereal-legume intercrop systems. European Journal of Agronomy doi: 10.1016/j.eja.2026.128065

Kottelenberg DB, Bastiaans L, Van Essen R, Kootstra G, Douma JC. 2026b. Can convolutional neural networks support agronomic analysis of cereal-legume canopy cover dynamics? Field Crops Research doi: 10.1016/j.fcr.2025.110236

Lafarge TA, Hammer GL. 2002. Predicting plant leaf area production: shoot assimilate accumulation and partitioning, and leaf area ratio, are stable for a wide range of sorghum population densities. Field Crops Research doi: 10.1016/S0378-4290(02)00085-0

Larue F, Fumey D, Rouan L, Soulie JC, Roques S, Beurier G, Luquet D. 2019. Modelling tiller growth and mortality as a sink-driven process using Ecomeristem: implications for biomass sorghum ideotyping. Annals of Botany doi: 10.1093/aob/mcz038

Li H, Jiang D, Wollenweber B, Dai T, Cao W. 2010. Effects of shading on morphology, physiology and grain yield of winter wheat. European Journal of Agronomy doi: 10.1016/j.eja.2010.07.002

Li C, Hoffland E, Kuyper TW, Yu Y, Zhang C, Li H, Zhang F, Van der Werf W. 2020. Syndromes of production in intercropping impact yield gains. Nature Plants doi: 10.1038/s41477-020-0680-9

Li S, van der Werf W, Zhu J, Guo Y, Li B, Ma Y, Evers JB. 2021. Estimating the contribution of plant traits to light partitioning in simultaneous maize/soybean intercropping. Journal of Experimental Botany doi: 10.1093/jxb/erab077

MacLaren C, Waswa W, Aliyu KT, Claessens L, Mead A, Schob C, Vanlauwe B, Storkey J. 2023. Predicting intercrop competition, facilitation, and productivity from simple functional traits. Field Crops Research doi: 10.1016/j.fcr.2023.108926

Montazeaud G, Violle C, Roumet P, Rocher A, Ecarnot M, Compan F, Maillet G, Fort F, Freville H. 2020. Multifaceted functional diversity for multifaceted crop yield: towards ecological assembly rules for varietal mixtures. Journal of Applied Ecology doi: 10.1111/1365-2664.13735

Morales A, Kottelenberg DB, Ernst A, Vezy R, Evers JB. 2025. The Virtual Plant Laboratory: a modern plant modeling framework in Julia. In Silico Plants doi: 10.1093/insilicoplants/diaf005

Morelli G, Ruberti I. 2000. Shade avoidance responses. Driving auxin along lateral routes. Plant Physiology doi: 10.1104/pp.122.3.621

Moss S. 2017. Herbicide Resistance in Weeds. In: Hatcher PE, Froud-Williams RJ, eds. John Wiley & Sons Ltd, 181–214.

Nielsen DC, Miceli-Garcia JJ, Lyon DJ. 2012. Canopy cover and leaf area index relationships for wheat, triticale, and corn. Agronomy Journal doi: 10.2134/agronj2012.0107n

Niinemets U. 2020. Leaf trait plasticity and evolution in different plant functional types. Annual Plant Reviews Online doi: 10.1002/9781119312994.apr0714

Oerke EC. 2006. Crop losses to pests. Journal of Agricultural Science doi: 10.1017/s0021859605005708

Panozzo A, Bolla PK, Barion G, Visioli G, Vamerali T. 2025. Morpho-physiological and agronomic responses of wheat varieties under artificial shade in agroforestry systems. Journal of the Science of Food and Agriculture doi: 10.1002/jsfa.70157

Perez GL, Torremorell A, Mugni H, et al. 2007. Effects of the herbicide roundup on freshwater microbial communities. Ecological Applications doi: 10.1890/07-0499.1

Poorter H, Niinemets U, Poorter L, Wright IJ, Villar R. 2009. Causes and consequences of variation in leaf mass per area (LMA): a meta-analysis. New Phytologist doi: 10.1111/j.1469-8137.2009.02830.x

R Core Team. 2025. R: a language and environment for statistical computing. Vienna, Austria: R Foundation for Statistical Computing. https://www.R-project.org/.

Salama HSA, Nawar AI, Khalil HE. 2022. Intercropping pattern and N fertilizer schedule affect the performance of additively intercropped maize and forage cowpea in the Mediterranean region. Agronomy doi: 10.3390/agronomy12010107

Sarlikioti V, De Visser PHB, Buck-Sorlin GH, Marcelis LFM. 2011. How plant architecture affects light absorption and photosynthesis in tomato: towards an ideotype for plant architecture using a functional-structural plant model. Annals of Botany doi: 10.1093/aob/mcr221

Schmitt J, Wulff R. 1993. Light spectral quality, phytochrome, and plant competition. Trends in Ecology and Evolution 8, 47–50.

Schmitt J, Stinchcombe JR, Heschel MS, Huber H. 2003. The adaptive evolution of plasticity: phytochrome-mediated shade avoidance responses. Integrative and Comparative Biology doi: 10.1093/icb/43.3.459

Schneider HM. 2022. Characterization, costs, cues and future perspectives of phenotypic plasticity. Annals of Botany doi: 10.1093/aob/mcac087

Smith H. 1982. Light quality, photoreception, and plant strategy. Annual Review of Plant Physiology 33, 481–518.

Sparkes DL, King M. 2008. Disentangling the effects of PAR and R:FR on lodging-associated characters of wheat (Triticum aestivum). Annals of Applied Biology doi: 10.1111/j.1744-7348.2007.00184.x

Spitters CJ, Toussaint HA, Goudriaan J. 1986. Separating the diffuse and direct component of global radiation and its implications for modeling canopy photosynthesis Part I. Components of incoming radiation. Agricultural and Forest Meteorology 38, 217–229.

Stomph T, Dordas C, Baranger A, et al. 2020. Chapter One - Designing intercrops for high yield, yield stability and efficient use of resources: are there principles? In: Donald LS, ed. Advances in Agronomy, vol 160. Academic Press, 1–50.

Van Bruggen AHC, He MM, Shin K, Mai V, Jeong KC, Finckh MR, Morris JG Jr. 2018. Environmental and health effects of the herbicide glyphosate. Science of the Total Environment doi: 10.1016/j.scitotenv.2017.10.309

Vandermeer J, Perfecto I. 2025. Inherent suboptimality of monocultural crop breeding. Agroecology and Sustainable Food Systems doi: 10.1080/21683565.2025.2475459

Vos J, Evers JB, Buck-Sorlin GH, Andrieu B, Chelle M, De Visser PHB. 2010. Functional-structural plant modelling: a new versatile tool in crop science. Journal of Experimental Botany doi: 10.1093/jxb/erp345

Wang R, Sun Z, Zhang L, et al. 2020. Border-row proportion determines strength of interspecific interactions and crop yields in maize/peanut strip intercropping. Field Crops Research doi: 10.1016/j.fcr.2020.107819

Wang W, Wang BZ, Zhang W, et al. 2026. Cereal-legume intercropping stabilizes yield and economic advantages under variable rainfall in semiarid rainfed environment. European Journal of Agronomy doi: 10.1016/j.eja.2025.127942

Yin X, Struik PC. 2009. C3 and C4 photosynthesis models: an overview from the perspective of crop modelling. NJAS: Wageningen Journal of Life Sciences doi: 10.1016/j.njas.2009.07.001

Zhu J, Van der Werf W, Anten NPR, Vos J, Evers JB. 2015. The contribution of phenotypic plasticity to complementary light capture in plant mixtures. New Phytologist doi: 10.1111/nph.13416

